# Computational metabolomics reveals overlooked chemodiversity of alkaloid scaffolds in *Piper fimbriulatum*

**DOI:** 10.1101/2024.12.10.627739

**Authors:** Tito Damiani, Joshua Smith, Téo Hebra, Milana Perković, Marijo Čičak, Alžběta Kadlecová, Vlastimil Rybka, Martin Dračínský, Tomáš Pluskal

## Abstract

Plant specialized metabolites play key roles in diverse physiological processes and ecological interactions. Identifying structurally novel metabolites, as well as discovering known compounds in new species, is often crucial for answering broader biological questions. The *Piper* genus (*Piperaceae* family) is known for its special phytochemistry and has been extensively studied over the past decades. Here, we investigated the alkaloid diversity of *Piper fimbriulatum*, a myrmecophytic plant native to Central America, using a metabolomics workflow that combines untargeted LC-MS/MS analysis with a range of recently-developed computational tools. Specifically, we leverage open MS/MS spectral libraries and metabolomics data repositories for metabolite annotation, guiding isolation efforts towards structurally-new compounds (i.e., dereplication). As a result, we identified several alkaloids belonging to 5 different classes and isolated one novel *seco*-benzylisoquinoline alkaloid featuring a linear quaternary amine moiety that we named fimbriulatumine. Notably, many of the identified compounds were never reported in Piperaceae plants. Our findings expand the known alkaloid diversity of this family, and demonstrate the value of revisiting well-studied plant families using state-of-the-art computational metabolomics workflows to uncover previously overlooked chemodiversity. To contextualize our findings into a broader biological context, we employed a workflow for automated mining of literature reports of the identified alkaloid scaffolds and mapped the results onto the angiosperm tree of life. By doing so, we highlight the remarkable alkaloid diversity within the *Piper* genus and provide a framework for generating hypotheses on the biosynthetic evolution of these specialized metabolites. Many of the computational tools and data resources used in this study remain underutilized within the plant science community. This manuscript demonstrates their potential through a practical application and aims to promote broader accessibility to untargeted metabolomics approaches.

**Significance Statement:** We combine untargeted metabolomics with a range of recently developed computational tools to uncover a previously overlooked diversity of alkaloid scaffolds in *Piper fimbriulatum*. Our findings demonstrate the potential of revisiting well-studied plant families using state-of-the-art computational metabolomics workflows to uncover previously overlooked chemodiversity.

## Introduction

Piperaceae (“pepper”) is a family of pan-tropical flowering plants and includes two of the largest genera of angiosperms, *Piper* (>2,400 species) and *Peperomia* (>1,400 species) (Simmonds et al. 2021). Numerous Piperaceae plants are of significant economic importance due to their widespread use as spices (e.g., black pepper) and in traditional medicine (e.g., Ayurveda) (Salehi et al. 2019). In particular, the *Piper* genus is known for its special phytochemistry and its phytochemical investigation over the past four decades has led to the isolation of over 300 different amide alkaloids (often referred to as “piperamides”) with a wide range of biological activities (Parmar et al. 1997; Martha Perez Gutierrez, Maria Neira Gonzalez, and Hoyo-Vadillo 2013; Salehi et al. 2019; Gómez-Calvario and Rios 2019). Despite such extensive exploration of the *Piper* phytochemistry, new piperamide structures are still regularly reported every year (Jung et al. 2024; Zhou et al. 2024). *Piper fimbriulatum* (**S. Figure 1**) is a myrmecophytic plant native to Central America that lives in symbiosis with *Pheidole bicornis* ants (Mayer, Schaber, and Hadacek 2008). Because of this symbiosis, the phytochemical characterization of this plant has focused mostly on its volatile compounds (terpenoids in particular), believed to play a role in the plant-insect communication (Mayer, Schaber, and Hadacek 2008; Mundina et al. 1998). During a plant metabolomics screening campaign (A. K. Jarmusch et al. 2022) we found evidence for *P. fimbriulatum* leaves to contain a high concentration of piperlongumine, a *Piper* alkaloid that has exhibited remarkable anticancer properties at a preclinical level (Conde et al. 2021). This finding, combined with the lack of reports of alkaloids in this plant, motivated a deeper investigation of the non-volatile chemodiversity of *P. fimbriulatum*, where the presence of unique specialized metabolites might have been overlooked.

**Figure 1.**
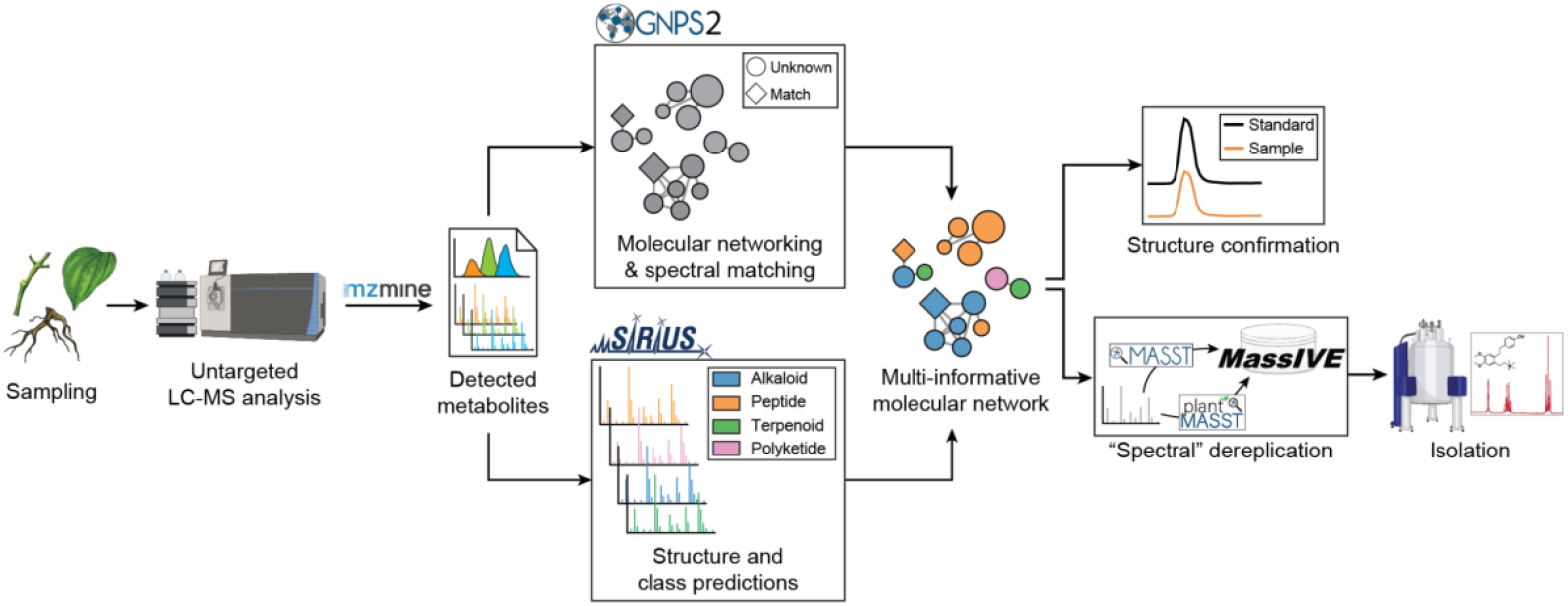
Analytical and computational workflow used in the present study. Different organs of the plant were sampled and analyzed by untargeted LC-MS/MS. Computational tools were used to aid the exploration of the detected chemical space, while publicly-available spectral libraries and data repositories were used to focus isolation efforts towards structurally-new phytochemicals.

Over the past two decades, liquid chromatography coupled to high-resolution tandem mass spectrometry (LC-MS/MS) has become the method of choice for metabolite profiling of crude plant extracts (Tsugawa et al. 2021). Such popularity is owed to high sensitivity and throughput of modern instruments, as well as the ability of tandem mass spectrometry data (i.e., MS/MS) to provide sufficient structural information for the annotation of the detected metabolites (S. A. Jarmusch et al. 2021). In addition, recently developed computational tools allow for efficient exploration of the metabolomics data and greatly assist the identification of new natural product analogs and scaffolds (Wolfender et al. 2019; Beniddir et al. 2021). Major advances in this direction were stimulated by the introduction of molecular networking (Watrous et al. 2012), which can capture structural relationships among (unknown) metabolites based on their fragmentation spectra (MS/MS or MS2), as well as the development of computational platforms to facilitate the (re)usability of open data (e.g., reference MS/MS libraries, metabolomics datasets) (M. Wang et al. 2016; Gomes et al. 2024). Finally, the recent establishment of open access resources for cataloging information available for known natural products (e.g., taxonomic occurrence, reported bioactivity), such as the LOTUS initiative (Rutz et al. 2022), allows for quick and reproducible mining of scientific literature, which can significantly aid data interpretation and dereplication (i.e., avoiding re-isolation of previously known compounds). Despite their potential, many of these computational tools remain underutilized within the plant science community, partly due to their recent introduction and, in our opinion, the perceived complexity for researchers outside the metabolomics field. The present manuscript showcases their application for the identification of specialized metabolites in non-model plant species, an often crucial task for addressing broader biological questions, and aims to promote broader accessibility to untargeted metabolomics approaches.

In this study, we used untargeted LC-MS/MS analysis to investigate the metabolite profiles of different organs of *P. fimbriulatum*. We primarily focused on alkaloids given the relevance of this compound class in the *Piper* genus. We used a range of computational tools to explore the (detected) chemodiversity of *P. fimbriulatum* and leveraged open MS/MS spectral libraries and metabolomics data repositories to direct our isolation efforts towards structurally-new compounds (**Figure 1**). We identified several alkaloids belonging to five different alkaloid classes, many of which have never been reported in Piperaceae. Moreover, we isolated one novel *seco*-benzylisoquinoline alkaloid with a linear quaternary amine moiety, which is an unusual moiety for plant alkaloids. Finally, we mined the Wikidata framework and compared the occurrence of these alkaloid classes across angiosperm orders and families. Our findings uncovered a remarkable diversity of alkaloid scaffolds being produced by the *Piper* genus, which goes beyond the well known piperamides. We believe that the workflows presented here can be applied to numerous biochemical and physiological studies requiring a deep and effective phytochemical characterization of specialized metabolism in non-model species.

## Results and discussion

### LC-MS analysis of *P. fimbriulatum*

In the present study, we used untargeted metabolomics to explore the chemodiversity of *P. fimbriulatum* (**S. Figure 1**) with a particular focus on alkaloids. After confirming the taxonomic identity of a collected specimen of *P. fimbriulatum* using DNA barcoding (see **Experimental Section**), we sampled the plant leaves, stems and root organs. From the collected samples, we prepared crude water-ethanol extracts and analyzed them using untargeted LC-MS/MS. The resulting raw data were processed using a pipeline of computational tools (**Figure 1**). We used the mzmine software for feature detection in order to convert the complex raw spectral data into a list of detected metabolites (i.e., LC-MS features) (Heuckeroth et al. 2024). Next, we used feature-based molecular networking (FBMN) (Nothias et al. 2020) to group together unknown metabolites with similar chemical structures based on the similarity between their MS/MS spectra. Finally, we annotated (unknown) metabolites in the obtained molecular network using a combination of MS/MS spectral matching, *in silico* prediction of chemical structures and compound classes using CSI:FingerID (Dührkop et al. 2015) and CANOPUS (Dührkop et al. 2021), with manual inspection of the prediction results (see **Metabolite annotation** section). FBMN resulted in the generation of 54 molecular networks (MNs) from 898 features (**Figure 2**), out of which only a small fraction (∼6%) could be annotated via direct match against the GNPS MS/MS spectral library. In order to aid the exploration of the detected chemical space, we layered the chemical structure and compound class predictions over the global molecular network (Mutabdžija et al. 2024). Overall, 245 nodes in the network were predicted as alkaloids and clustered into 5 main molecular networks (**Figure 2**, hereafter referred to as **MN1, MN2, MN3, MN4, MN5**), which we focused our annotation efforts on.

**Figure 2.**
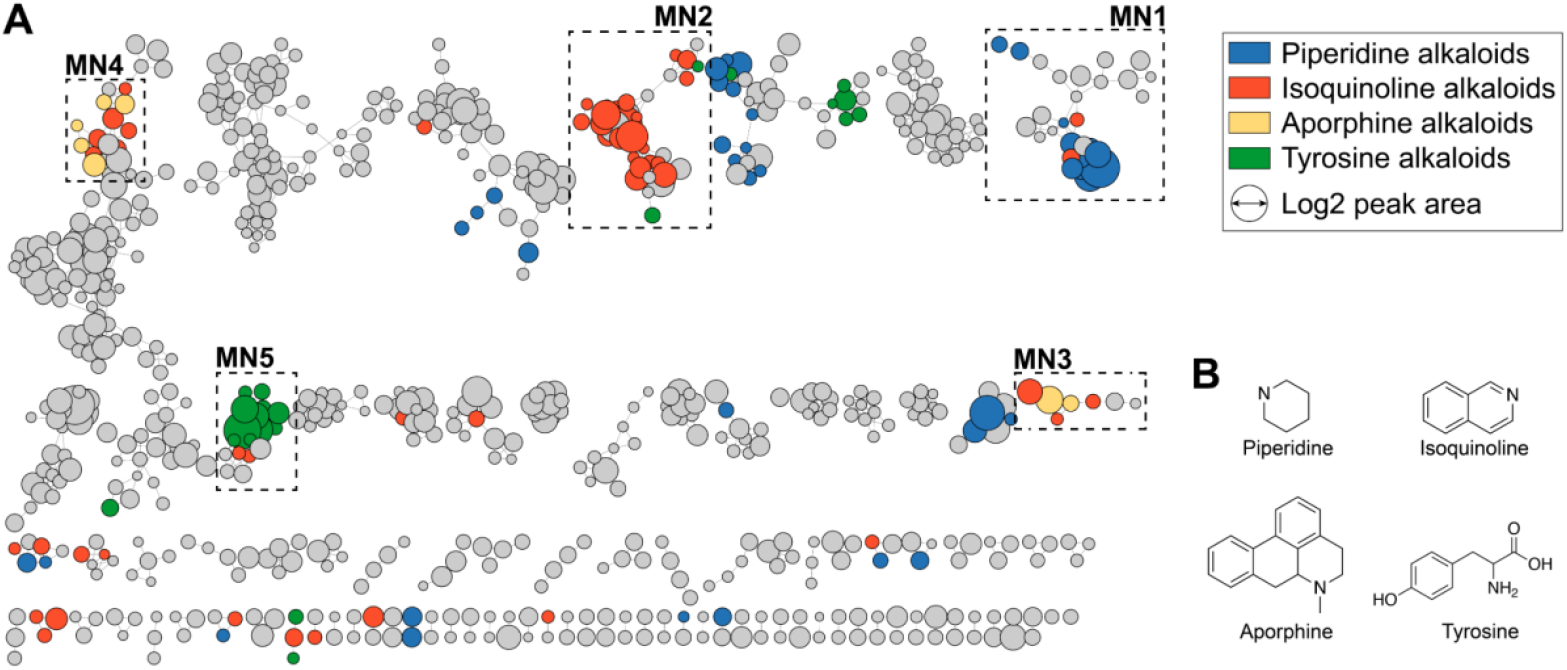
A) A global molecular network of the chemical space detected in *P. fimbriulatum* (leaf, stem and root organs). Nodes predicted as alkaloids by CANOPUS are colored based on the alkaloid class. Alkaloid-related molecular families are highlighted as MN1-5. B) Chemical scaffolds corresponding to the alkaloid classes highlighted in the global molecular network.

### Piperidine alkaloids molecular network (MN1)

MN1 contained 29 nodes, out of which 12 were predicted as “piperidine alkaloids” (**Figure 2A**), a class of lysine-derived alkaloids characterized by the presence of a piperidine ring (**Figure 2B**) in their chemical structures (Szőke, Lemberkovics, and Kursinszki 2013). Spectral matching against the GNPS MS/MS library retrieved a highly-confident hit (cosine similarity >0.96, **S. Figure 2A**) to piperlongumine (**1**), in accordance with our screening campaign results (see **Introduction**). Moreover, several other spectra in MN1 were annotated as piperlongumine analogs (i.e., a similar MS/MS spectrum, but different precursor *m/z*) with high confidence (cosine similarity >0.93). We confirmed the annotation of piperlongumine using a commercial standard (**S. Figure 3A**) and used this confirmation as a starting point to “propagate” the annotation throughout MN1 (**S. Note 1**, see also **Metabolite annotation** section). By doing so, we putatively annotated 5 piperlongumine analogs (**2**), (**3**), (**4**), (**5**), (**6**); see **S. Table 1**), and one homodimer (**7**). Next, we synthesized pure standards (see **Experimental Section**) for compounds (**2**), (**3**), (**5**), (**6**) and confirmed our annotations by retention time and MS/MS spectral matching (**S. Figure 3B-E**). Regarding compound (**7**), [2+2] cycloaddition reactions are known to occur under visible and UV light (Hurtley, Lu, and Yoon 2014; Nguyen and Al-Mourabit 2016). Therefore, we generated a mixture of piperlongumine dimers by irradiation of the pure monomer with 365 nm UV light (see **Experimental section**). LC-MS/MS analysis of the reaction mixture confirmed the presence of (**7**) in the native plant (**S. Note 1, S. Figure 7A-B**). Interestingly, we observed a nearly-identical chromatographic profile between the reaction mixture and the leaf extract (**S. Figure 7C**). This suggests that the formation of these dimers in the plant might be UV-induced, rather than enzyme-catalyzed (**S. Note 1**). Finally, spectral matching retrieved a highly-confident hit (cosine similarity >0.99, **S. Figure 2B**) to piperine (**8**), the main alkaloid in *Piper nigrum* fruits, which we also confirmed using a commercial standard (**S. Figure 3F**). Interestingly, piperine was detected exclusively in the root organ, unlike the other piperidine alkaloids (**Figure 3C**). Piperine and other other piperamides were recently reported to localize within specific cells of the fruit perisperm and the root cortex in *P. nigrum*.(Jäckel et al. 2022) Given that *P. fimbriulatum* belongs to the same genus, it is reasonable to expect a similar phenomenon. However, our data allow us to assess only the inter-organ distribution, as the analysis was performed on bulk samples (i.e., leaf, stem, root). Investigating inter-tissue and cellular localization of these alkaloids would require spatial metabolomics techniques with tissue-or cellular-level resolution (e.g., single-cell metabolomics or laser microdissection), which we plan to pursue in future studies.

**Figure 3.**
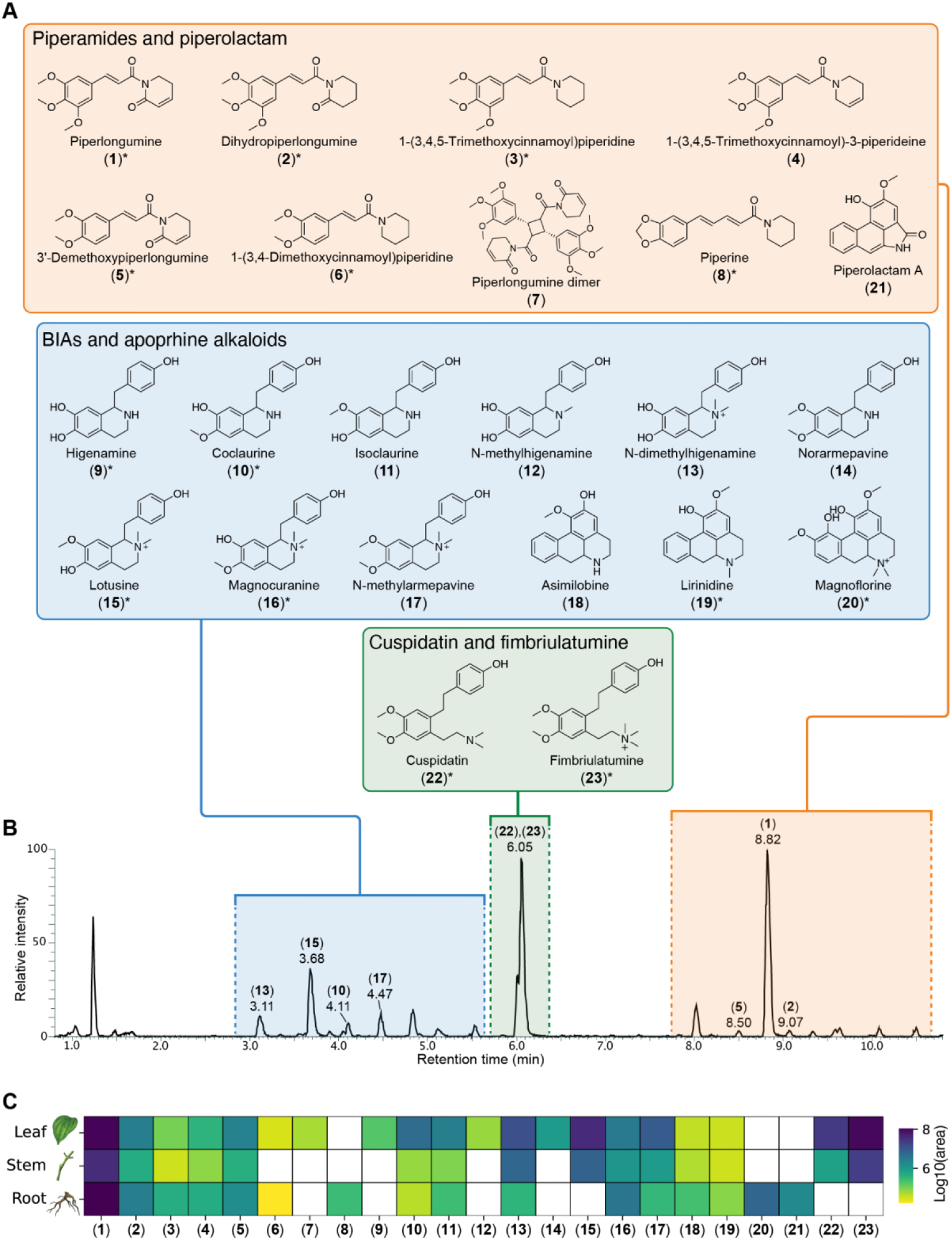
A) Chemical structures of the *P. fimbriulatum* alkaloids identified in the present study. Compounds confirmed by retention time match with commercial standard or NMR structural characterization are marked with asterisks. More information about the identified compounds are provided in **S. Table 1**; B) Base peak LC-MS chromatogram of *P. fimbriulatum* leaf acquired in positive ionization mode; C) Heatmap representing the abundance (shown as log10-transformed LC-MS peak area) of the annotated alkaloids across the different plant organs (i.e., leaf, stem, root). It must be noted that the heatmap aims at highlighting the inter-organ distribution of individual metabolites. Since absolute quantification for each compound was not performed, abundance comparisons between metabolites (e.g., compound (**1**) more abundant in the plant than compound (**15**)) cannot be made due to differences in ionization efficiency and other factors.

### Isoquinoline alkaloids molecular network (MN2)

MN2 contained 34 nodes, out of which 12 were predicted as “isoquinoline alkaloids”, which are a large and structurally-diverse group of specialized metabolites (∼2,500 known structures) predominantly found in the plant kingdom (Yang et al. 2024). Isoquinoline alkaloids are further classified in benzylisoquinolines (BIAs), aporphines and other subclasses based on their structural backbone, and are known to possess potent pharmacological properties (e.g., the narcotic morphine and the antimicrobial berberine) (Yang et al. 2024). Spectral matching against the GNPS MS/MS library retrieved highly-confident hits (cosine similarity >0.95) to higenamine (**9**) and coclaurine (**10**) (**S. Figure 2C-D**), two central intermediates in the biosynthesis of all plant BIAs (Tian et al. 2024). We confirmed both these annotations using commercial standards (**S. Figure 8A-B**) and, similar to what was done for MN1, we used these confirmations to “propagate” the annotation throughout MN2. Overall, we annotated 9 BIAs (**S. Table 1**), including *N*-methylarmepavine (**17**) and its intermediates, and confirmed 4 of these annotations using commercial standards (**S. Figure 8A-D, S. Note 2**). Overall, BIAs showed higher accumulation in the leaves, unlike piperamides which were more often found consistently across all organs (**Figure 3C**).

### Aporphine and aristolactams alkaloids molecular networks (MN3 and MN4)

MN3 and MN4 contained various nodes predicted as either “isoquinoline” or “aporphine alkaloids” by CANOPUS. Aporphine alkaloids are a major class of BIAs, including more than 700 reported compounds (da Silva Mendes et al. 2023). Many aporphine alkaloids have shown promising therapeutic effects, particularly for the treatment of central nervous system diseases (Carbone et al. 2019) and metabolic syndromes (F.-X. Wang et al. 2021). Spectral matching against the GNPS MS/MS library retrieved hits to asimilobine (**18**), magnoflorine (**20**), and piperolactam A (**21**) (**S. Figure 2E-G**). Piperolactams (a.k.a. aristolactams) are a relatively small group of phytochemicals with a variety of reported pharmacological activities, mainly occurring in plants from the Annonaceae, Aristolochiaceae and Piperaceae families (Kumar et al. 2004). Similar to what was done for MN1 and MN2, we used commercial standards (when available) and manual inspection of the MS/MS spectra to confirm these annotations (**S. Note 3**). Overall, we confirmed the annotation for lirinidine (**19**) and magnoflorine (**20**) using commercial standards (**S. Figure 8E-F**), and putatively annotated asimilobine (**18**) and piperolactam A (**21**) based on MS/MS spectral matching against the GNPS MS/MS library and manual inspection of the results. Interestingly, all these alkaloids were almost exclusively detected in the root organ of the plant (**Figure 3C**). Such organ-specific production may result from the localized expression of biosynthetic enzymes, which could be leveraged in future studies to elucidate the biosynthetic pathway(s) for these alkaloids. In fact, organ-specific metabolite abundance can help narrow down candidate genes for experimental validation, focusing on those with similar expression patterns (see, for example, (Schnabel et al. 2021)).

### Fimbriulatumine alkaloids molecular network (MN5)

Finally, MN5 contained 18 nodes, out of which 13 were predicted as “tyrosine alkaloids” by CANOPUS. Interestingly, two nodes in the network (*m/z* 330.207 and 344.223, respectively) corresponded to dominant peaks in the LC-MS chromatogram of *P. fimbriulatum* leaves (see **Figure 3B**). No reliable match against the GNPS MS/MS spectral library was retrieved for any node within the network, thereby complicating the annotation process. For both peaks, CSI:FingerID provided low-confidence (score <0.3) chemical structure predictions and the manual inspection of the MS/MS spectra did not reveal fragment peaks or patterns common to other identified metabolites, which suggested these compounds could belong to a distinct alkaloid class. Before moving to the purification and structural characterization of these metabolites, we used an MS/MS-based dereplication strategy (see **Figure 1** and **Experimental Section**) to reduce the chance of (re-)isolating already-known compounds. We started by using the recently-developed plantMASST (Gomes et al. 2024) tool to verify whether a similar MS/MS spectrum was ever collected in publicly-available plant metabolomics datasets. In particular, plantMASST searches MS/MS spectra across a curated database of LC-MS/MS data from plant extracts within the GNPS ecosystem which, at the time of writing, covers more than 2’700 species from 1’400 genera and 246 botanical families (Gomes et al. 2024). The plantMASST search revealed that both metabolites were exclusively found in *P. fimbriulatum* (**S. Figure 12** and **S. Figure 13**). Next, we searched the same MS/MS spectra against a wider range of public metabolomics data repositories using the Mass Spectrometry Search Tool (MASST) (M. Wang et al. 2020; Mongia et al. 2024). While plantMASST limits the search to curated plant LC-MS/MS datasets, MASST enables querying of the entire GNPS/MassIVE(M. Wang et al. 2016) data repository (containing over 16’000 datasets and 8 billion of spectra at the time of writing). For both peaks, MASST returned no match outside datasets deposited by our own lab (MASST search 1, MASST search 2). Given the potential structural novelty, we proceeded with the targeted isolation of these compounds from the original plant material (see **Experimental section**). NMR analysis of the purified compounds confirmed the structure of two *seco*-benzyltetrahydroisoquinolines alkaloids (**22**) and (**23**) (**S. Figure 21-31**), respectively carrying a tertiary and quaternary linear amine moiety (**Figure 3**). While we found a previous report of (**22**), named cuspidatin, from the bark of *Antidesma cuspidatum (Elya et al*., *n*.*d*.), a tropical plant from the Phyllanthaceae family, we did not find previous report for (**23**) neither in primary literature, nor in chemical structure databases (i.e., PubChem, Reaxys, SciFinder). Given the structural novelty and its unique occurrence in the *P. fimbriulatum* species (see **S. Figure 13**), we named the compound fimbriulatumine.

### Distribution of *Piper* alkaloids in angiosperms

All identified alkaloids in this study are shown in **Figure 3A** and summarized in **S. Table 1**. While *Piper* plants are mainly known for the production of piperamides (amide alkaloids), here we confirmed the occurrence of several non-amide alkaloids in *P. fimbriulatum*. Surprised by such diversity of alkaloid structures and scaffolds in a single *Piper* species, we contextualized our findings in an evolutionary framework using the recently-published angiosperms tree of life(Zuntini et al. 2024), which covers 58% of the approximately 13,600 currently accepted genera and arguably constitutes the most comprehensive phylogenomic reconstruction of this clade’s evolution at the time of writing. **Figure 4** shows the angiosperm phylogenetic tree mapped with literature reports, mined using Wikidata (see **Experimental section** and **S. Figure 14**), of the alkaloid scaffolds we identified in *P. fimbriulatum* (i.e., benzylisoquinoline, aporphine, piperolactam, piperidine, *seco*-benzylisoquinoline). We found the widest distribution for the BIAs (203 genera from 48 families and 24 orders) and aporphine alkaloids (141 genera from 23 families and 16 orders) scaffolds. Within the Piperaceae family, we found only two reports of BIAs and aporphine alkaloids: one in *Piper argyrophylum* dating back 1996 (Singh et al. 1996) and, more recently, a second in *Piper sarmentosum(Ware et al*. *2024)*. Our work suggests a potential wider occurrence of these alkaloids within the *Piper* genus. Concerning the piperolactam scaffold, we found reports in 14 genera from 6 families and 3 orders and, interestingly, it always co-occurs with BIAs and/or aporphine alkaloids, except for two genera in the Saururaceae family (Zhuang et al. 2014; Z. Wu et al. 2021). This supports the hypothesis that piperolactams are derived from 4,5-dioxoaporphine via benzilic acid rearrangement(Kumar et al. 2004), although evidence for this is still lacking. In contrast to the wide occurrence of the structural cores described above, we found the piperidine scaffold to be reported (almost) exclusively in Piperaceae plants, with only one report outside this family -i.e., *Punica granatum (Chaturvedi et al*. *2013)* (Lythraceae family, Myrtales order). Finally, we found reports for the *seco*-benzylisoquinoline scaffold only in *Polyalthia insignis (Lee, Chuah, and Goh 1997)* (Annonaceae family, Magnoliales order) and *Antidesma cuspidatum* (Phyllanthaceae family, Malpighiales order), while we report it for the first time in the *Piper* genus in this study. Notably, *Piper* is the only genus across the phylogenetic tree where all of the 5 considered scaffolds were reported, highlighting its remarkable alkaloid diversity.

**Figure 4.**
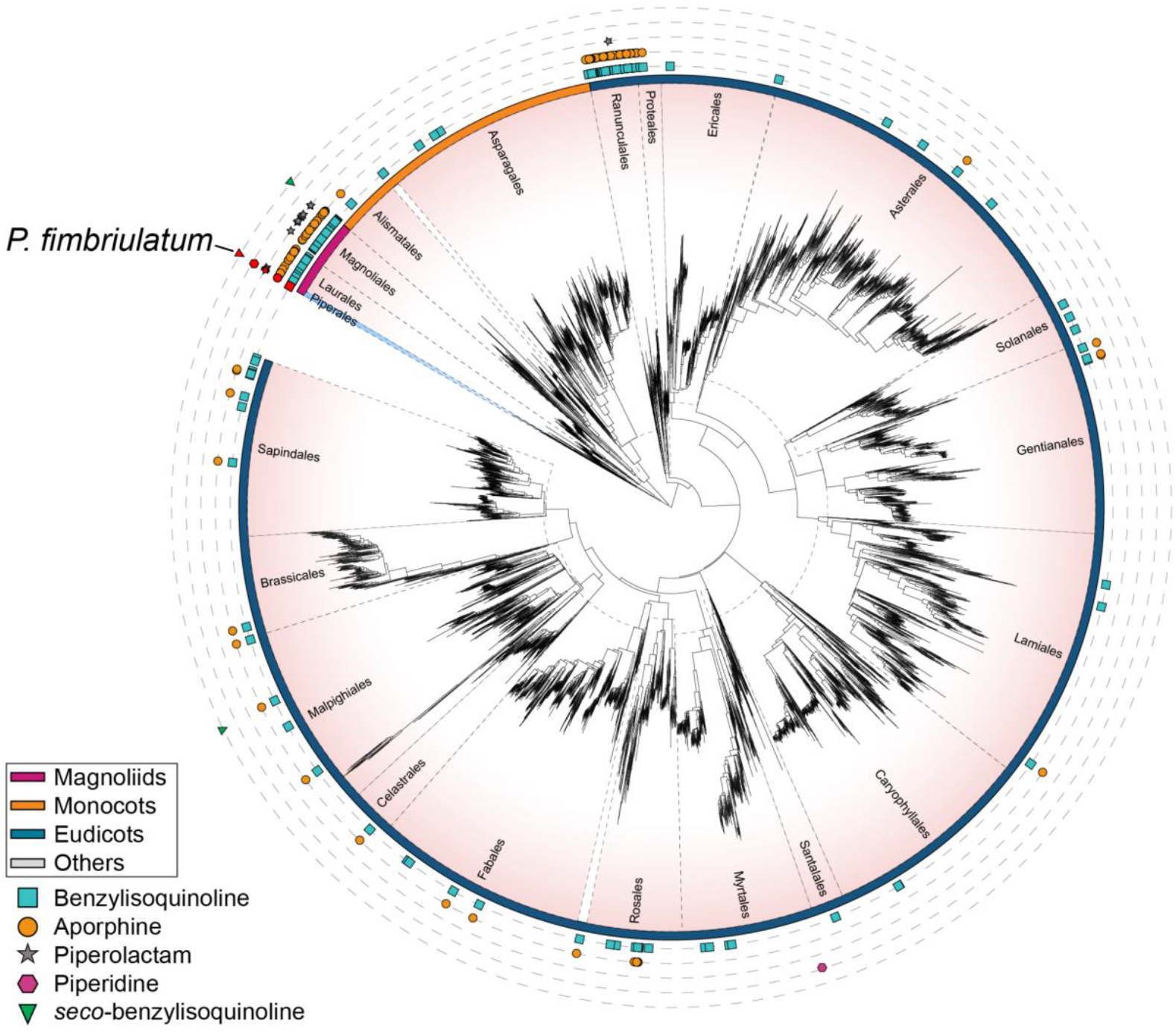
Angiosperm tree of life from (Zuntini et al. 2024) mapped with literature reports, mined from Wikidata, of the alkaloid scaffolds considered in the present study (i.e., benzylisoquinoline, aporphine, piperolactam, piperidine, *seco*-benzylisoquinoline). Each leaf in the tree corresponds to a representative species for each genus as described in the original publication. Different plant orders are separated by dashed black lines and the Piperales order is highlighted in light blue. Reports for each scaffold are represented with different colored shapes (see legend in the figure). Reports for the *Piper* genus are highlighted in red. Colored arcs around the tree indicate the four main clades of angiosperms as described in the original publication: Magnoliids, Monocots, Eudicots. The tree in the figure only shows the plant orders for which at least one scaffold was reported. A full version of the tree is provided in **S. Figure 15**. More details about the construction of the tree are provided in the **Experimental section**. The figure was created using iTOL(Letunic and Bork 2024). The original tree can be accessed at https://itol.embl.de/tree/14723112167224931731658296.

We believe that the workflow shown in **S. Figure 14** provides a powerful approach for efficiently mining knowledge bases (e.g., Wikidata) and tracking the occurrence of NPs across the plant kingdom, which, in our opinion, remains largely underestimated. This can be crucial for studies addressing broader biological questions, such as uncovering the evolutionary history of these specialized metabolic pathways, by enabling the generation of new hypotheses. For instance, the presence of the benzylisoquinoline and aporphine scaffolds in the early-diverging Piperales order may support their monophyletic evolution in angiosperms.(Liscombe et al. 2005) In contrast, the occurrence of the piperidine scaffold exclusively in the *Piper* and *Punica* genera might indicate convergent evolution of piperidine alkaloid biosynthesis, considering the large phylogenetic distance between these two genera. While proving this claim would require a much deeper investigation (see, for example, (Huang et al. 2016)) we postulate that leveraging knowledge bases combined with insightful visualization strategies can significantly support this exploration.

## Conclusions

In the present study, we investigated the alkaloid diversity of *P. fimbriulatum* using untargeted LC-MS/MS metabolomics. We used a range of computational tools to assist the exploration of the detected chemical space and leveraged open MS/MS spectral libraries and data repositories to direct our isolation efforts towards structurally-novel compounds. Overall, we documented the natural occurrence of 23 alkaloids belonging to 5 different classes, including a novel *seco*-benzylisoquinoline alkaloid carrying a linear quaternary amine nitrogen, which we named fimbriulatumine. Next, we contextualized these findings within the angiosperm evolutionary framework by mining the Wikidata open knowledge base and mapping the results onto the angiosperm tree of life. This analysis highlighted a remarkable diversity of alkaloid scaffolds within the *Piper* genus.

While expanding the known alkaloid diversity of the Piperaceae family and confirming it as a great source of specialized metabolites, our findings demonstrate the value of revisiting well-studied plant families using state-of-the-art computational metabolomics tools to uncover overlooked chemodiversity. In fact, many of the identified alkaloids were never reported in Piperaceae plants, despite their extensive phytochemical investigation over the past four decades. Another highlight of our work is the effectiveness of the “spectral dereplication” in the proposed LC-MS/MS data analysis workflow, which leverages the ever-growing MS/MS spectral libraries and metabolomics data repositories to efficiently annotate both known and structurally novel metabolites. This is demonstrated by the fact that, apart from the new fimbriulatumine, all other alkaloid structures were confirmed simply by purchasing the corresponding analytical standard, thus saving significant time and resources that would have otherwise been spent on the (re)isolation of known compounds.

We believe that the analytical and computational workflows presented here can be instrumental in improving our capabilities for the phytochemical characterization of non-model plant species. This is crucial for studies addressing broader biological questions, such as physiological processes and ecological interactions, which often rely on identifying novel metabolites mediating these processes. Moreover, a more effective identification and reporting of structurally-novel metabolites, as well as known compounds in new species, will accelerate efforts to map their distribution across the plant kingdom, which, in our opinion, remains largely underestimated. This could ultimately lead to the generation of new evolutionary hypotheses about the biosynthetic origin of specialized metabolic pathways.

Despite their potential, many of the computational tools employed in this study have not yet become routine resources for many researchers outside the metabolomics field. For this reason, we ensured the full reproducibility of the computations and results by sharing all generated data and scripts in open repositories. We hope this will help non-experts familiarize themselves with the computational tools used in this manuscript and facilitate their broader adoption within the plant science community.

## Experimental Section

### Chemicals

Commercial analytical standards Piperlongumine (>97% purity), piperidine (99% purity) and higenamine (95% purity) were purchased from Merck-Sigma-Aldrich (Prague, Czech Republic). Higenamine, coclaurine, lotusine, magnocurarine, lirinidine, and magnoflorine were all obtained in a natural product compound library from Targetmol (Wellesley, USA). Chemical synthesis compounds: 3,4,5-Trimethoxy cinnamic acid (>98% purity) was purchased from BLDpharm (Turnov, Czech Republic). Dichloromethane (99.8% purity) and Tetrahydrofuran (99.5% purity) extra dry over molecular sieve from Fisher Scientific (Germany). Pivaloyl-chloride (99% Purity), Triethylamine (> 99.5% purity) 2-Piperidinone were from Merck-Sigma-Aldrich (Prague, Czech Republic). Solvents and additives for LC-MS analysis (i.e., acetonitrile, water, formic acid, all Optima LCMS Grade) were purchased from Fisher Scientific (Germany). Concerning the molecular biology reagents, ultra-pure agarose was purchased from Invitrogen-Fisher (Prague, Czech Republic). NaCl p.a. from Penta Chemicals (Prague, Czech Republic), molecular biology-grade NaHCO_3_, Na_2_SO_4_, MgSO_4_ and CTAB extraction buffer from SERVA (Heidelberg, Germany). DMSO molecular biology grade from Merck-Sigma-Aldrich (Prague, Czech Republic).

### Sample collection and taxonomic confirmation

*P. fimbriulatum* samples were collected from the Prague Botanical Garden in January 2021 (specimen n° 2016.00648). Leaf, stem and root organs were collected in triplicate from a single plant. After collection, samples were immediately stored in dry ice and transferred to −80 °C until analysis. The taxonomic identity of *P. fimbriulatum* was confirmed through DNA barcoding on the internal transcribed spacer (ITS) region, which proved to be an efficient marker for *Piper* species differentiation (Jaramillo et al. 2008). Genomic DNA was isolated from 50 mg of fresh leaf tissue using the cetyltrimethylammonium bromide (CTAB) protocol as described in Aboul-Maaty et al. (Aboul-Maaty and Oraby 2019). The ITS region was amplified by polymerase-chain reaction (PCR) using the Phusion High-Fidelity DNA Polymerase (M0530L, New England Biolabs) and the following primers (as reported in (Jaramillo et al. 2008)):

- ITS-A (forward, 5’→3’): GGAAGGAGAAGTCGTAACAAGG
- ITS-B (reverse, 5’→3’): CTTTTCCTCCGCTTATTGATATG

PCR reactions were conducted in a 3 × 32-well ProFlex thermocycler (Thermo Fisher) and in 20 μL volume: 12.75 µL of nuclease-free water, 4 µL of 5X HF buffer, 1 μL of 10 μM forward primer (ITS-A), 1 μL of 10 μM reverse primer (ITS-B), 0.5 µL of 10 mM dNTPs mix, 0.5 µL of template DNA, 0.25 µL of Phusion DNA polymerase. After an initial denaturation step at 98 °C for 30 seconds, the following thermocycling conditions were applied for 35 cycles: 98 °C for 10 seconds (denaturation), 61 °C for 15 seconds (annealing), 72 °C for 30 seconds (extension). The reaction was concluded with a final extension step at 72 °C for 10 min. PCR products were separated by gel electrophoresis (80 V for 45 minutes) on a 1% (w/v) agarose gels and purified using NucleoSpin silica gel cartridges (Macherey-Nagel) prior to sequencing.

Sanger sequencing was performed and a consensus sequence was generated by aligning the sequencing results from both primers. The resulting consensus sequence was searched against the NCBI database using the BLAST alignment tool (Camacho et al. 2009). The top match (100% identity, 99% query coverage, E-value of 0) was retrieved against an ITS sequence deposited for *P. fimbriulatum* (GenBank EF056251.1; NCBI:txid425151). Raw sequencing results and the resulting consensus sequence are provided in the **Supplementary information**.

### Metabolite extraction

Frozen plant material was homogenized under liquid nitrogen using mortar and pestle. 50 mg (±5 mg) of material was weighed in a pre-frozen 2 mL plastic tube with round bottom. Samples were further homogenized by adding a pre-frozen stainless steel bead to each tube and high-speed shaking (30 seconds, 25 rpm) using TissueLyser II (QIAGEN). For secondary metabolites extraction, each sample was added with 1 mL of ethanol/H_2_O (75:25) solution, incubated for 5 minutes at 40 °C, and shaken again for 60 seconds at 25 rpm. We opted for this extraction mixture as, based on preliminary testing, it offered the best compromise in terms of metabolite coverage and (relative) extraction yield. Following the extraction step, samples were centrifuged for 10 minutes at 14000 rpm, 750 μL of supernatant was transferred into new 1.5 mL tubes and dried using a SpeedVac vacuum concentrator (Thermo Fisher). Finally, samples were resuspended in 750 μL of H_2_O/CH_3_CN (50:50) solution and transferred to glass vials for LC-MS analysis. Extraction blanks were prepared and analyzed in order to identify and remove signals arising from contaminants introduced during the extraction process (see **Feature detection** section).

It is important to note that we have not performed absolute quantification of the metabolites reported in this manuscript (see **Results** section). Therefore, the relative abundances of detected metabolites may be influenced by their ionization efficiency, the chosen extraction mixture, as well as the resuspension in H_2_O/CH_3_CN (50:50) prior to LC-MS analysis. For this reason, abundance comparisons between metabolites (see, for example, **Figure 3**) should be made with this in mind.

### LC-MS analysis

LC-MS/MS analyses were performed using a Vanquish UHPLC system (Thermo Fisher Scientific) coupled to an Orbitrap ID-X mass spectrometer equipped with a heated electrospray ionization (HESI) source. Reverse-phase (RP) separation of analytes was performed on an Acquity BEH C18 column, 150 mm x 2.1 mm, 1.7 μm particle size (Waters). The temperature in the autosampler and column oven was set to 10 °C and 40 °C, respectively. The mobile phase consisted of water (A) and CH_3_CN (B), both containing 0.1% formic acid. A constant flow rate of 350 μL/min was used and the chromatographic gradient was as follow: initial hold of 0.5 min at 5% of B, linear gradient of 15 min from 5 to 100% of B, isocratic step of 1.8 min at 100% of B for column wash, isocratic step of 2 min at 5% of B for column reconditioning. The injection volume was set to 1 μL. HESI source parameters were set to 50 arbitrary units (AU) of sheath gas, 12 AU of auxiliary gas, 1 AU of sweep gas, vaporizer temperature of 350 °C, spray voltage of 3000 V, ion transfer tube temperature of 325 °C. MS data were acquired in positive ionization and data dependent acquisition mode (DDA), with MS2 spectra collected with a cycle time of 0.6 seconds (i.e., maximum time between MS1 scans). MS1 data were collected in profile mode, 100–1000 *m/z* scan range, 60’000 resolution, 1 microscan, 45% RF lens, normalized AGC target of 50% (i.e., ≈2E5), maximum ion injection time of 118 milliseconds. MS2 data were collected in profile mode, 15’000 resolution, 1 microscan, normalized AGC target of 100% (i.e., ≈5E4), maximum ion injection time of 80 milliseconds. Selected precursor ions were fragmented with a quadrupole isolation window of 0.8 *m/z*, 1E5 minimum intensity threshold, dynamic exclusion of 2 seconds (5 ppm tolerance, isotope exclusion set to ‘ON’), fixed normalized HCD collision energy of 35%, apex detection set to ‘ON’ with expected peak width (FWHM) of 4 seconds and desired apex window of 50%. Raw LC-MS/MS data were visualized using Freestyle v1.8 (Thermo Fisher Scientific).

### LC-MS data processing

#### Feature detection and feature-based molecular networking

Raw LC-MS data were converted from Thermo Fisher proprietary format (.RAW) to open format (.mzML) using the MSConvert software (*ProteoWizard*) (Chambers et al. 2012). Untargeted feature detection was performed using the mzmine software (Schmid et al. 2023) (v4.2.0) as described in Heuckeroth et al. (Heuckeroth et al. 2024). Briefly, the mass detection noise level was set to 5.0E4 and 2.0E3 for MS1 and MS2 level, respectively, and all signals below these intensities were discarded. Extracted ion chromatograms (XIC) were built for each *m/z* with a 0.002 Da or (5 ppm) tolerance and retained if exhibiting at least 7 consecutive data points above 1.5E5 intensity, and a minimum absolute height or 5.0E5. After smoothing (LOESS method, 4-scans window), XICs were resolved (Local Minimum Resolver module) using the following parameters: chromatographic threshold=90%; minimum search range=0.05; minimum absolute height=5.0E5; minimum ratio of peak top/edge=1.70; peak duration range=0.0-2.0 minutes; minimum scans=5. MS2 spectra were paired to the resolved peaks with a 0.002 Da or (5 ppm) “MS1 to MS2 precursor” tolerance and using feature edges as retention time (RT) limits. Redundant ^13^C isotope features were filtered using 0.001 Da (or 3.5 ppm) and 0.01 minutes tolerances; a monoisotopic shape of the isotopic pattern was required and the most intense isotope was kept as representative. Feature alignment (Join aligner module) was performed using 0.002 Da (or 5 ppm) and 0.08 minutes sample-to-sample tolerances (equal weight was attributed to m/z and RT). Duplicated features (i.e., potential artifacts from the alignment) were removed using 0.0005 Da (or 2 ppm) and 0.02 minutes tolerances. Features detected in the extraction blank samples were removed from the aligned feature table unless they exhibited a 3x fold-change compared to the blank. Correlation grouping and ion identity networking were carried out using defaults parameters (see provided batch file). Finally, features that were not detected in at least 3 samples (since each plant organ was collected and analyzed in triplicate) and/or did not contain at least 2 isotopes in the isotopic pattern were filtered out. The resulting feature table was exported for further analysis by FBMN and the SIRIUS software. For FBMN, MS/MS spectra were merged across samples (weighted average) using the following parameters: intensity merge mode=sum; expected mass deviation: 0.02 Da (or 5 ppm); cosine threshold=0; signal count threshold=0; isolation window offset=0; isolation window width of 0.8 Da. For SIRIUS, MS/MS spectra were not merged and exported with a *m/z* tolerance of 0.002 Da (or 5 ppm). The above-described feature detection steps and parameters are also provided as a mzmine configuration batch file (see **Supplementary Information**).

FBMN was performed through the Global Natural Products Social Molecular Networking (GNPS) platform (M. Wang et al. 2016) using the following parameters: 0.01 Da parent mass tolerance, 0.01 Da MS/MS fragment ion tolerance, minimum cosine similarity of 0.7, minimum 6 matched peaks. The FBMN results were visualized using Cytoscape (v3.10.2) (Shannon et al. 2003).

#### Metabolite annotation

For the (putative) annotation of unknown metabolites in the molecular network, we used a combination of MS/MS spectral library matching (Bittremieux, Wang, and Dorrestein 2022), *in silico* prediction of chemical structures and compound classes (Dührkop et al. 2015, 2021) and manual interpretation of MS/MS spectra. During the FBMN workflow in GNPS, all spectra are automatically searched against the public reference MS/MS libraries hosted on the platform. Both exact and analog matches with cosine similarity above 0.7 and ≥6 matched fragment peaks were retrieved. “Analog matches” are retrieved by using a spectral similarity score that takes into account both fragment peaks and neutral losses in the similarity calculation. To increase the annotation coverage, we computationally annotated molecular formulas, chemical structures and compound classes for all MS/MS spectra in the network using the SIRIUS software (v5.8.5) (Dührkop et al. 2019). Molecular formulas were computed with the `formula` module by matching the experimental and predicted isotopic patterns (Böcker et al. 2009) and from fragmentation tree analysis (Böcker and Dührkop 2016) of MS/MS. Default parameters were used, except for: instrument type=orbitrap; mass accuracy for MS1=5 ppm; mass accuracy for MS2=7.5 ppm. Formula predictions were refined with the ZODIAC module (Ludwig et al. 2020) using default parameters. *In silico* structure annotation was done with CSI:FingerID (Dührkop et al. 2015) using structures from biological databases. Systematic compound class annotations were obtained with CANOPUS (Dührkop et al. 2021) and using default parameters and the NPClassifier (Kim et al. 2021) ontology.

For the MS/MS-based dereplication, we used MASST (M. Wang et al. 2020; Mongia et al. 2024) and plantMASST (Gomes et al. 2024). MASST (Mass Spectrometry Search Tool) is a MS/MS similarity search tool for querying MS/MS spectra of unknown molecules against public data repositories, such as the GNPS/MassIVE repository (over 16’000 metabolomics datasets and 8 billion of spectra at the time of writing). Spectral matches are returned along with dataset(s) metadata, which can provide biological context information about the queried MS/MS spectrum, even when the corresponding chemical structure is unknown. PlantMASST is a domain-specific version of MASST, which searches the query MS/MS spectrum within a curated reference database of LC-MS/MS data acquired from plant extracts. Spectral matches are returned along with metadata information about the plant species, genus, etc. At the time of writing, the plantMASST reference database contains data from over 19’000 plant extracts covering 246 botanical families, 1’400 genera, and 2’700 species (Gomes et al. 2024). Both MASST and plantMASST searches were performed with the following default parameters: precursor mass tolerance=0.05 Da; fragment mass tolerance=0.05 Da; cosine threshold=0.7; minimum matched peaks=3; analog search=OFF.

Manual interpretation of MS/MS spectra was done using a range of dedicated software tools, such as mzmine, SIRIUS, Metabolomics Spectrum Resolver (Bittremieux et al. 2020) and the GNPS Dashboard (Petras et al. 2022). In particular, to propagate the annotation from confirmed metabolites to neighboring nodes in the molecular network, we took advantage of the recently-developed ModiFinder (Shahneh et al. 2024) tool, which compares the MS/MS spectra of a known structure and its unknown analog to locate the most likely modification site. All the putative annotations were manually inspected and, whenever possible, confirmed with either commercially-available and *in-house* synthetized analytical standards (see **Chemical synthesis of piperamides** section)

### Chemical synthesis of piperamides

Compounds (**2**), (**3**), (**5**) and (**6**) were synthetized as described in (Y. Wu et al. 2014) and (Rao et al. 2012) (**S. Figure 16A-B**). For compounds (**2**) and (**5**), in argon atmosphere, trimethoxy cinnamic acid (1.0 eq) was dissolved in dry THF (5 mL), cooled to 0 °C and triethylamine (1.5 eq) was added. Pivaloyl chloride (1.1 eq) was added dropwise in the reaction mixture and stirred for 1h at 0 °C. Afterwards, 2-piperidinone (1.1 eq) was dissolved in dry THF (2 mL), cooled to −78 °C and 1.6 M *n*-BuLi in hexane (1.1 eq) was added dropwise and stirred for 1h at −78 °C. Previously prepared mixture of anhydride was added to 2-piperidinone reaction mixture dropwise at −78 °C and stirred for 1h. The reaction mixture was quenched with saturated NH_4_Cl (5 mL), diluted with H_2_O (30 mL) and extracted with EtOAc (3×25 mL). Combined organic layers were washed with saturated NaCl (30 mL), dried over anhydrous MgSO_4_ and evaporated to dryness. Crude product was purified by RP-flash chromatography (50 g column, H_2_O/MeOH, 0–100%), yielding pure (**2**) and (**5**). For compounds (**3**) and (**6**), trimethoxy cinnamic acid (1.00 eq) was dissolved in dry DCM (5 mL), cooled to 0 °C and then triethylamine (2 eq) was added. Pivaloyl chloride (1.1 eq) was added dropwise in the reaction mixture and stirred for 1h at 0 °C. Afterwards, piperidine (1.5 eq) was added dropwise and stirred overnight. Triethylamine (1.100 eq) was added to a stirred solution of piperidine (1.5 eq) in dry DCM (2 mL) at 0 °C and stirred for 1h. The reaction mixture was added dropwise to anhydride reaction mixture at 0 °C and stirred overnight. Reaction mixture was diluted with DCM, washed with saturated NaHCO_3_ (20 mL) and saturated NaCl (20 mL), dried over Na_2_SO_4_, concentrated and purified by RP-flash chromatography (50 g column, H_2_O/MeOH, 0– 100%), yielding pure (**3**) and (**6**). All pure compounds were dissolved in DMSO-*d*_6_ for the subsequent NMR measurements. The NMR spectra were measured on a 400-MHz Bruker Bruker Avance III HD spectrometer. Chemical shifts are reported in ppm (δ) relative to solvent (DMSO-*d*_6_) signal: δ = 2.54 and 40.45 ppm for ^1^H and ^13^C, respectively (**S. Figure 17-20**).

For the synthesis of piperlongumine dimer, piperlongumine (**1**) standard was dissolved in 1.67 mL of methanol (2 mM), transferred to a glass vial, and evaporated in the dark under gentle nitrogen flow. This resulted in thin, homogeneous film of the compound. Film was then irradiated with a UV lamp at 365 nm., At 0, 1, 5, 10 and 30 minutes irradiation was stopped, film was dissolved in MeOH as above, a time point was taken (10 μL), and then again evaporated into a film. After irradiation, samples were immediately stored in the dark, at −20^°^C until analysis. Aliquots were diluted to 1 μM concentration and analyzed by LC-MS.

### Isolation of cuspidatin and fimbriulatumine

Lyophilized, ground leaves (≈2 g) of *P. fimbriulatum* were sequentially extracted by maceration four times with 150 mL of methanol for 1.5 hours (each round), at room temperature and protected from direct light sources. The resulting solutions were filtered and combined in a round bottom flask and evaporated under reduced pressure at 40 °C, yielding ≈500 mg of a dark-green residue. The obtained extract was redissolved in 20 mL of methanol and transferred to a separatory funnel and washed three times with 10 mL of *n*-octane. The methanol fractions resulting from each wash were pooled and evaporated again, yielding ≈350 mg of residue. The so-obtained residue was dissolved in methanol (concentration ≈100 mg/mL), centrifuged and and fractionated using a preparative HPLC system (1290 Infinity II Preparative Binary Pump, Agilent Technologies) equipped with a UV-Vis detector (1260 Infinity II Variable Wavelength Detector, Agilent Technologies) and fraction collector (1290 Infinity II Preparative Open-Bed Fraction Collector, Agilent Technologies). Sample was separated by reversed-phase chromatography using a XBridge C18 column, 10 mm × 250 mm, 5 μm particle size (Waters). The temperature in the column oven was set to 40 °C. The mobile phase consisted of water (A) and MeOH (B), both containing 0.1% formic acid. A constant flow rate of 17 mL/min was used and the chromatographic gradient was as follow: initial hold of 3 min at 10% of B, linear gradient of 62 min from 10 to 70% of B, isocratic step of 5 min at 100% of B for column wash, isocratic step of 5 min at 10% of B for column reconditioning. The injection volume was set to 200 μL and the sample was injected multiple times until collection of sufficient material for structural confirmation via NMR. This separation yielded ≈1 mg of compounds (**22**) and (**23**). All compounds were dissolved in CD_3_CN for subsequent structural confirmation by NMR. The NMR spectra were measured on a 500-or a 600-MHz Bruker Avance III HD spectrometer (^1^H at 500.0 or 600.1 MHz and ^13^C at 125.7 or 150.9 MHz). The spectra were referenced to solvent signals (*δ* = 1.94 and 1.32 ppm for ^1^H and ^13^C, respectively). The signal assignment was performed using a combination of 1D (^1^H and ^13^C) experiment and 2D correlation experiments (H,H-COSY, H,C-HSQC and H,C-HMBC). In some cases, the sample quantity was insufficient for the detection of some or all ^13^C signals in the 1D experiments. The structural analysis and signal assignment was then based on the 2D correlation experiments. Through literature search, we found a previous report (Elya et al., n.d.) of compound (**22**), with NMR data deposited in the SpectraBase repository (SpectraBase ID: HqNww9TxQ60).

### Mapping alkaloid scaffolds to the phylogeny of angiosperms

To investigate the distribution of specific alkaloid scaffolds across angiosperms, we designed SPARQL queries to retrieve natural products containing one of the 5 alkaloid scaffolds reported in *P. fimbriulatum* in the present study (i.e., benzylisoquinoline, aporphine, piperolactam, piperidine, *seco*-benzylisoquinoline, **S. Figure 14**). The queries targeted reported natural product structures containing the scaffold (sub)structure (defined using SMILES string), along with the corresponding plant species they were isolated from, as documented in Wikidata based on literature reports. The original SPARQL queries are available in the **Supplementary Information** and in the GitHub repository (see **Data availability** section). Query results were manually inspected and refined as follows. First, structures that were retrieved by the initial query but clearly unrelated to the target scaffold were filtered out. This was achieved by defining chemical substructures common to the “unwanted” structure retrieved by the initial query (**S. Figure 14B**) and using the `HasSubstructMatch` module from the RDKit Python library (v2023.03.2). The substructures used for this filtering are shown in **S. Figure 14C** and **D**, and both unfiltered and filtered datasets are included in the **Supplementary Information**. In addition, to facilitate review and reproducibility of this manuscript, we created a Jupyter Notebook (`clean_wikidata.ipynb`, available in the GitHub repository) that interactively visualizes the substructures removed for each scaffold. After structural filtering, we manually inspected the cleaned datasets to identify false or unreliable reports due to errors in Wikidata and/or primary literature. In particular, the query results for the piperidine scaffold included erroneous reports of piperine in *Capsicum* species (attributed to an error in Wikidata; Weaver et al., 1984) and entries for *Centaurea aegyptica (Dahmy et al*. *1985)* and *Aglaia perviridis (Zhang et al*. *2010)*, both excluded due to lack of NMR data in the original publication. Next, we mapped the so-cleaned literature reports onto the recently published phylogenomics tree of angiosperms (Zuntini et al. 2024). In particular, the tree file named `global_tree_brlen_pruned_renamed.tree` was used (see (Rizzo Zuntini and Carruthers 2024)). The latter contains one representative species per genus, therefore, the mapping was done at the genus-level - i.e., the genus names were extracted from the literature reports and matched with the genera present in the angiosperm tree. This was done using custom Python scripts (`03_clean_wikidata.py` and `05_create_itol_annotation.py`) available in the GitHub repository. Phylogenetic tree visualization and figure creation was done using Interactive Tree Of Life (iTOL) online tool (Letunic and Bork 2024). The phylogenetic tree displayed in **Figure 4** is a smaller version of the original tree created by removing plant orders for which no report for any alkaloid scaffold was retrieved (see Python scripts `05_create_small_tree.py` in the GitHub repository). The tree can be accessed at https://itol.embl.de/tree/14723112167224931731658296. The full tree is shown in **S. Figure 15** and can be accessed at https://itol.embl.de/tree/14723112167277531728383616.

## Data availability

Raw LC-MS files (.mzML), mzmine configuration batch file and processing results are available through Zenodo (https://zenodo.org/records/14336729). NMR data are also available through Zenodo at the same link. SPARQL queries and corresponding output (both raw and filtered), phylogenetic trees and iTOL annotation files are also available through Zenodo at the same link. All code and software is available through GitHub under the following link https://github.com/pluskal-lab/PiperFIM. The FBMN results are available at the following link: https://gnps2.org/status?task=43853471d153488b95144abd3af36a9d. Annotated phylogenetic trees can be accessed at https://itol.embl.de/tree/14723112167277531728383616 (**Figure 4**), and https://itol.embl.de/tree/14723112167224931731658296 (**S. Figure 15**).

## Acknowledgments

We thank Klára Lorencová, Petr Vacík and Ludvík Bortl from the Prague Botanical Garden for their support with the sample collection. T.P. was supported by the Czech Science Foundation (GA CR) grant 21-11563M and by the European Union’s Horizon 2020 research and innovation programme under Marie Skłodowska-Curie grant agreement No. 891397. T.D. was supported by the European Regional Development Fund; P JAC; Project “IOCB MSCA PF Mobility” (No. CZ.02.01.01/00/22_010/0002733). T.H. was supported by the European Union’s Horizon Europe research and innovation program under the Marie Skłodowska-Curie grant agreement No. 101130799. M.P. received support from the Visegrad Fund for presenting this research at conferences.

