## Supplementary Material for "Computational metabolomics reveals overlooked chemodiversity of alkaloid scaffolds in *Piper fimbriulatum*"

### Supplementary Tables

**S. Table 1**. List of *P. fimbriulatum* alkaloids identified in the present study.

| Name | Adduct | *m/z* | RT (min) | MZmine ID | Annotation |
|---|---|---|---|---|---|
| Piperlongumine (**1**) | $[M+H]^+$ | 318.1342 | 8.83 | 452 | Standard |
| Dihydropiperlongumine (**2**) | $[M+H]^+$ | 320.1497 | 9.07 | 463 | Standard |
| 1-(3,4,5-Trimethoxycinnamoyl)piperidine (**3**) | $[M+H]^+$ | 306.1705 | 8.90 | 458 | Standard |
| 1-(3,4,5-Trimethoxycinnamoyl)-3-piperideine (**4**) | $[M+H]^+$ | 304.1548 | 9.56 | 503 | MS/MS |
| 3'-Demethoxypiperlongumine (**5**) | $[M+H]^+$ | 288.1235 | 8.50 | 406 | Standard |
| 1-(3,4-Dimethoxycinnamoyl)piperidine (**6**) | $[M+H]^+$ | 276.1598 | 8.58 | 411 | Standard |
| Piperlongumine dimer (**7**) | $[M+H]^+$ | 635.2605 | 10.41 | 561 | MS/MS |
| Piperine (**8**) | $[M+H]^+$ | 286.1442 | 10.12 | 555 | Standard |
| Higenamine (**9**) | $[M+H]^+$ | 272.1284 | 3.31 | 130 | Standard |
| Coclaurine (**10**) | $[M+H]^+$ | 286.1442 | 4.11 | 202 | Standard |
| Isococlaurine (**11**) | $[M+H]^+$ | 286.1442 | 5.11 | 270 | MS/MS |
| N-methylhigenamine (**12**) | $[M+H]^+$ | 286.1441 | 3.88 | 185 | MS/MS |
| N-dimethylhigenamine (**13**) | $[M]^+$ | 300.1599 | 3.11 | 123 | MS/MS |
| Norarmepavine (**14**) | $[M+H]^+$ | 300.1599 | 4.91 | 254 | MS/MS |
| Lotusine (**15**) | $[M]^+$ | 314.1756 | 3.68 | 161 | Standard |
| Magnocurarine (**16**) | $[M]^+$ | 314.1755 | 3.90 | 186 | Standard |
| N-methylarmepavine (**17**) | $[M]^+$ | 328.1912 | 4.47 | 227 | MS/MS |
| Asimilobine (**18**) | $[M+H]^+$ | 268.1335 | 5.32 | 279 | MS/MS |
| Lirinidine (**19**) | $[M+H]^+$ | 282.1494 | 5.83 | 298 | Standard |
| Magnoflorine (**20**) | $[M]^+$ | 342.1706 | 4.41 | 225 | Standard |
| Piperolactam A (**21**) | $[M+H]^+$ | 266.0813 | 8.39 | 403 | MS/MS |
| Cuspidatin (**22**) | $[M+H]^+$ | 330.2067 | 6.02 | 309 | NMR |
| Fimbriulatumine (**23**) | $[M]^+$ | 344.2226 | 6.06 | 315 | NMR |

### Supplementary Figures

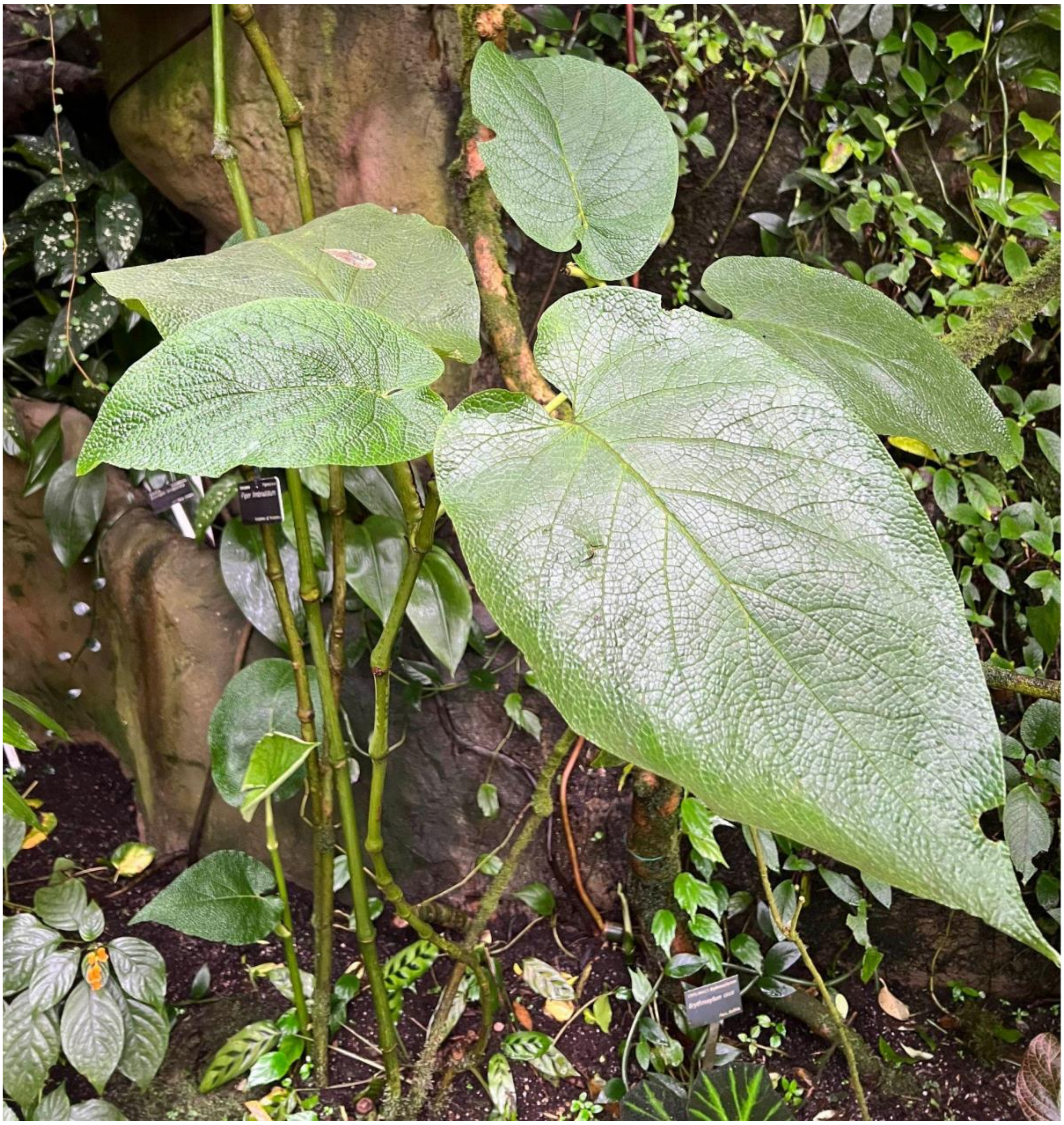

**S. Figure 1**. Photo of the *Piper fimbriulatum* plant sampled in Prague Botanical Garden.

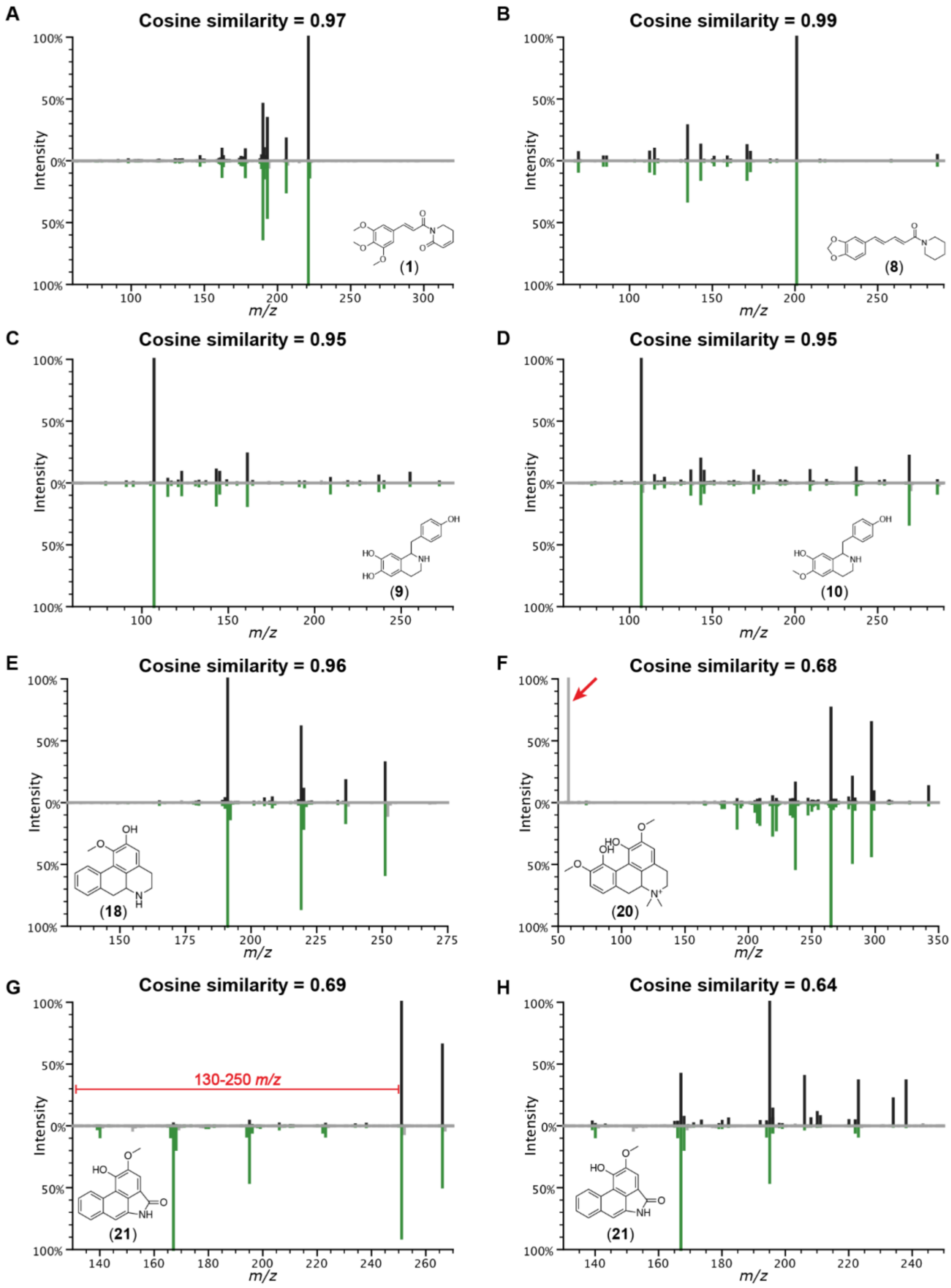


**S. Figure 2**. Spectral matches against the GNPS MS/MS spectral library. Aligned fragment peaks are highlighted in black for the experimental spectrum (top) and green for the library spectrum (bottom). Links to reproduce the MS/MS mirror plots in the figure with the Metabolomics Spectrum Resolver are provided. A) Piperlongumine (**1**) (CCMSLIB00004720841), link; B) Piperine (**8**) (CCMSLIB00005761602), link; C) Higenamine (**9**) (CCMSLIB00006439446), link; D) Coclaurine (**10**) (CCMSLIB00005436045), link; E) Asimilobine (**18**) (CCMSLIB00005436078), link; F) Magnoflorine (**20**) (CCMSLIB00010127350), link. The red arrow indicates the fragment peak *m/z* 58.065 in the experimental spectrum that was not matched due to the fact that the library spectrum was acquired with acquisition range starting from *m/z* 60; G) Piperolactam A (**21**) (CCMSLIB00000851803), link. The *m/z* range 130-250 (highlighted in red) is shown in more detail in the following panel; H) Piperolactam A (**21**), magnification of *m/z* range 130-250 of the spectrum shown in the previous panel, link.

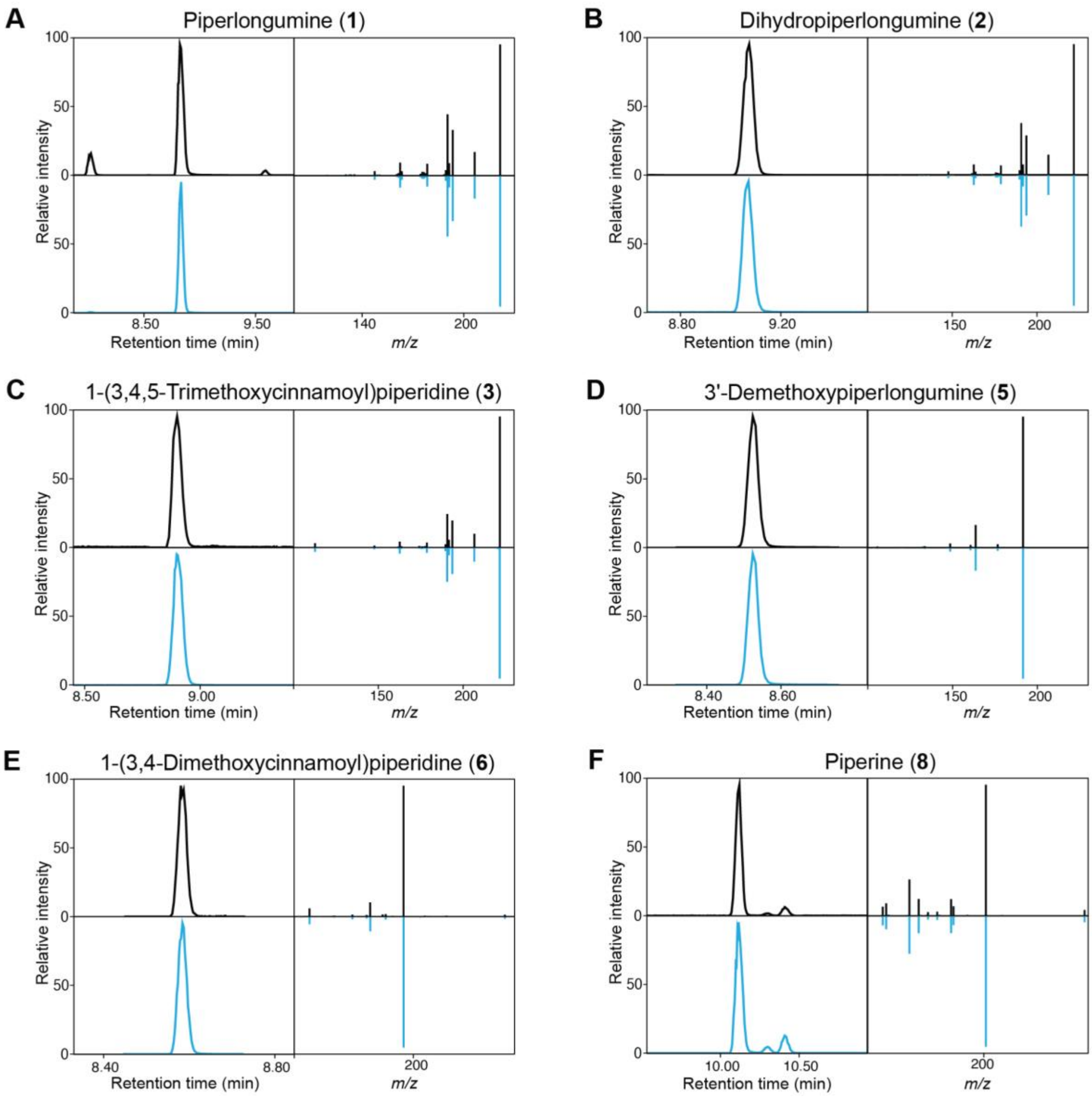


**S. Figure 3**. Confirmation of piperamide annotations via retention time matching with analytical standards. Extracted ion chromatograms and MS/MS spectra of sample (top) and standards (bottom) are coloured in black and blue, respectively. A) Piperlongumine (**1**), *m/z* 318.134; B) Dihydropiperlongumine (**2**), *m/z* 320.149; C) 1-(3,4,5-Trimethoxycinnamoyl)piperidine (**3**), *m/z* 306.170; D) 3'-Demethoxypiperlongumine (**5**), *m/z* 288.123; E) 1-(3,4-Dimethoxycinnamoyl)piperidine (**6**), *m/z* 276.159; F) Piperine (**8**), *m/z* 286.143.

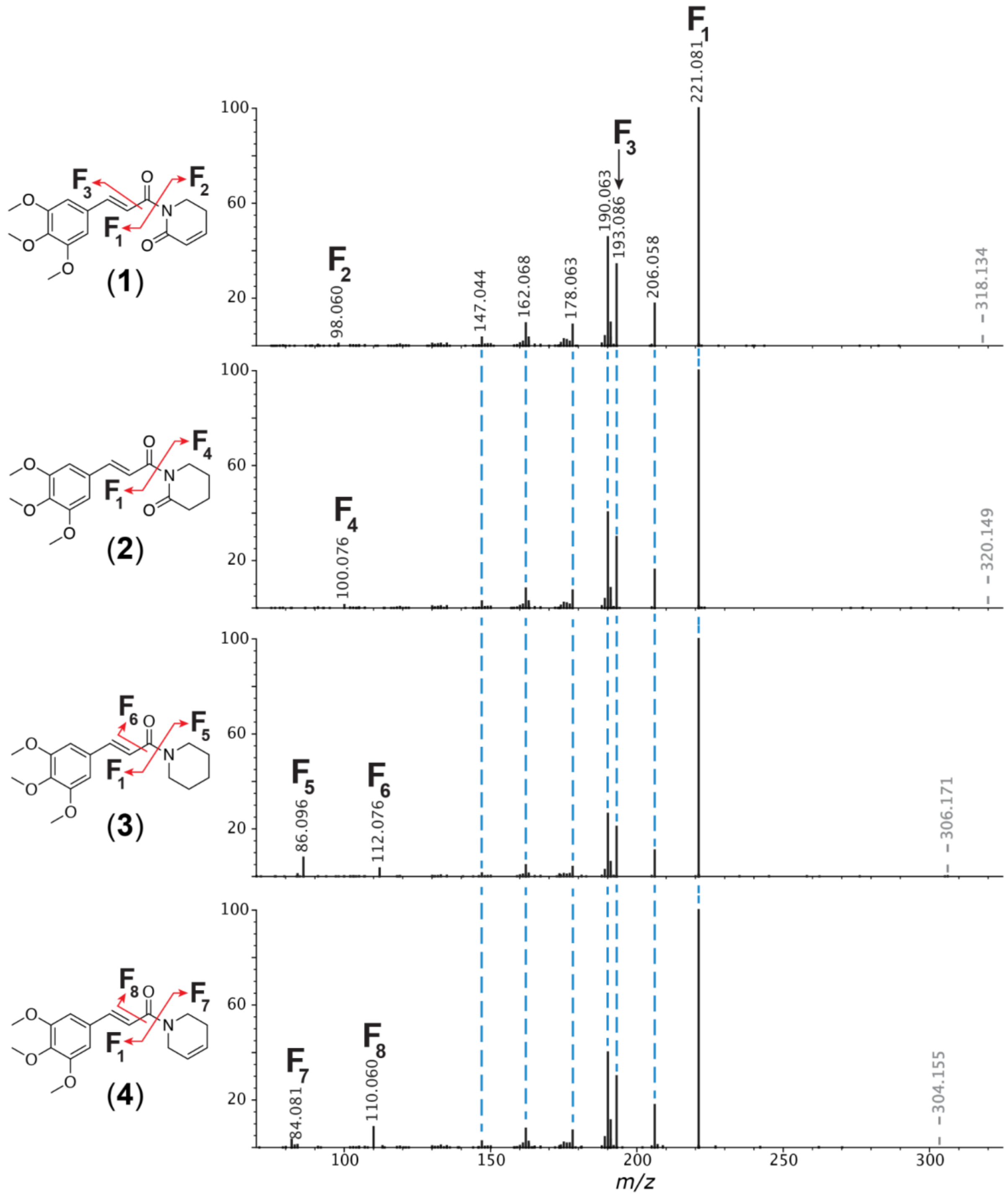


**S. Figure 4**. Interpretation of piperamides MS/MS spectra. Dashed gray lines represent precursor *m/z*. Dashed blue lines represent matching fragment peaks. Proposed fragmentation pathways are represented by red arrows in the chemical structures. MS/MS spectra in the figure can be reproduced using the Metabolomics Spectrum Resolver at the following links: Piperlongumine, Dihydropiperlongumine, 1-(3,4,5-Trimethoxycinnamoyl)piperidine; 1-(3,4,5-Trimethoxycinnamoyl)-3-piperideine.

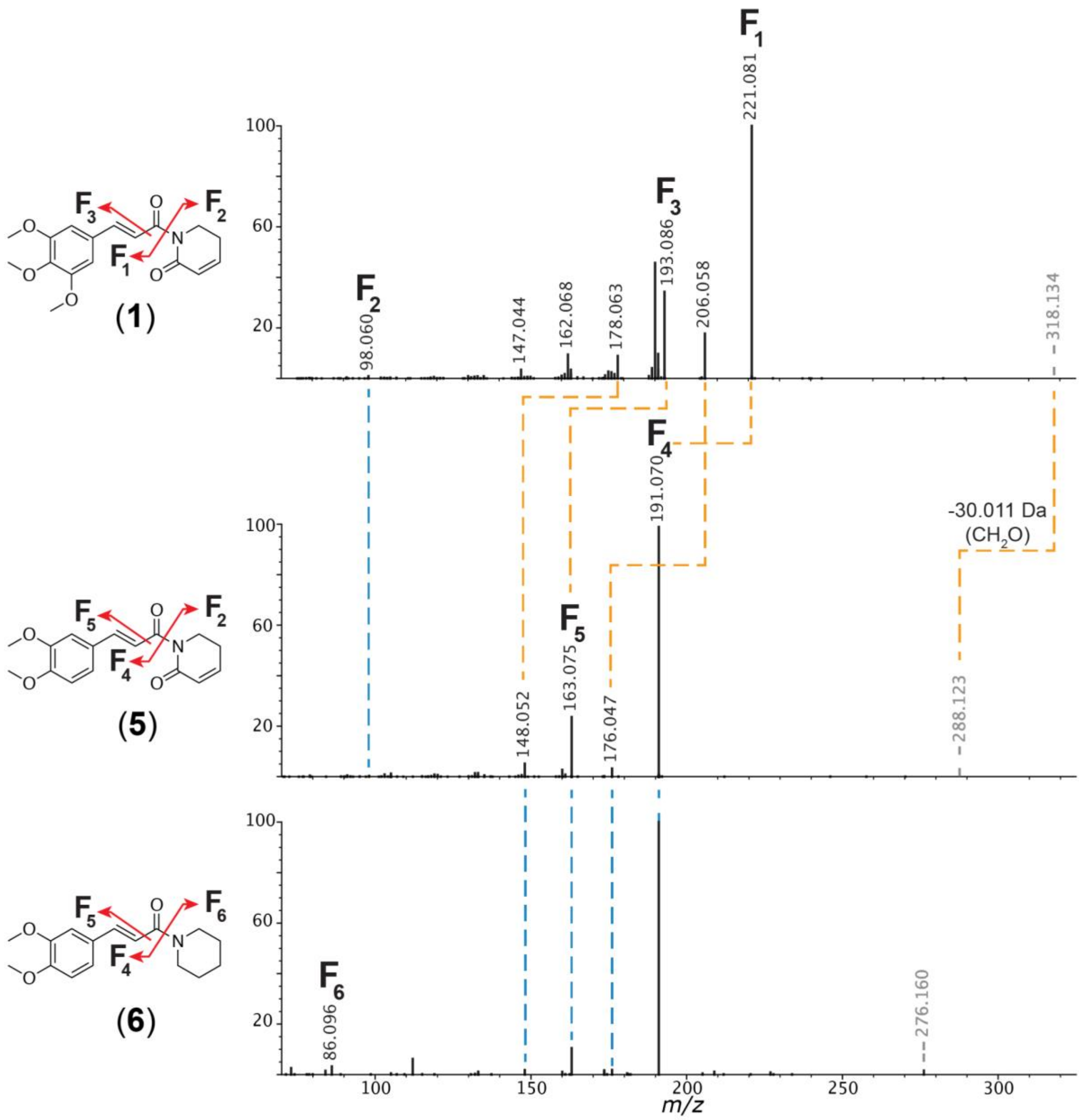


**S. Figure 5**. Interpretation of piperamides MS/MS spectra. Dashed gray lines represent precursor *m/z*. Dashed blue lines represent matching fragment peaks. Dashed orange lines represent matching fragment peaks shifted by the same difference as the precursors' *m/z* (-30.011 Da). Proposed fragmentation pathways are represented by red arrows in the chemical structures. MS/MS spectra in the figure can be reproduced using the Metabolomics Spectrum Resolver at the following links: piperlongumine, 3'-Demethoxypiperlongumine, 1-(3,4-Dimethoxycinnamoyl)piperidine.

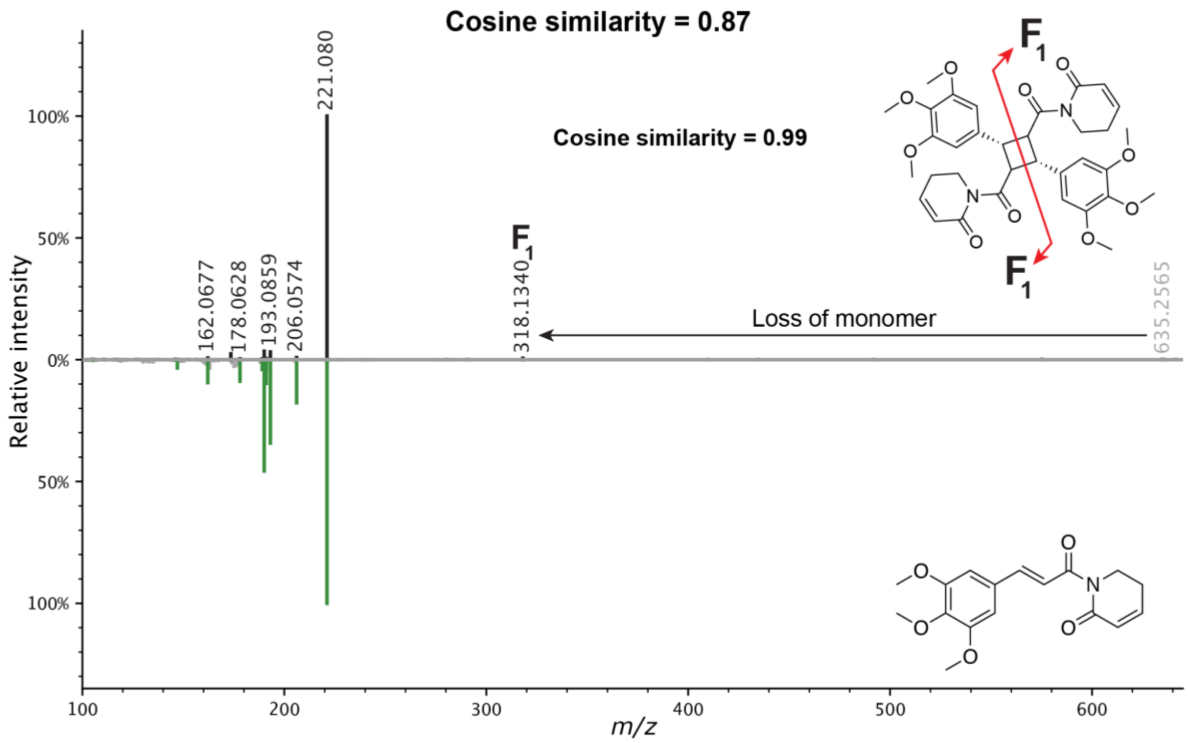


**S. Figure 6**. MS/MS mirror plot of piplartin dimer (**7**) and piperlongumine (**1**). Precursor *m/z* is highlighted in gray. Aligned fragment peaks are highlighted in black for the piplartin dimer spectrum (top) and green for the piperlongumine spectrum (bottom). The MS/MS mirror plot in the figure can be reproduced using the Metabolomics Spectrum Resolver at this link.

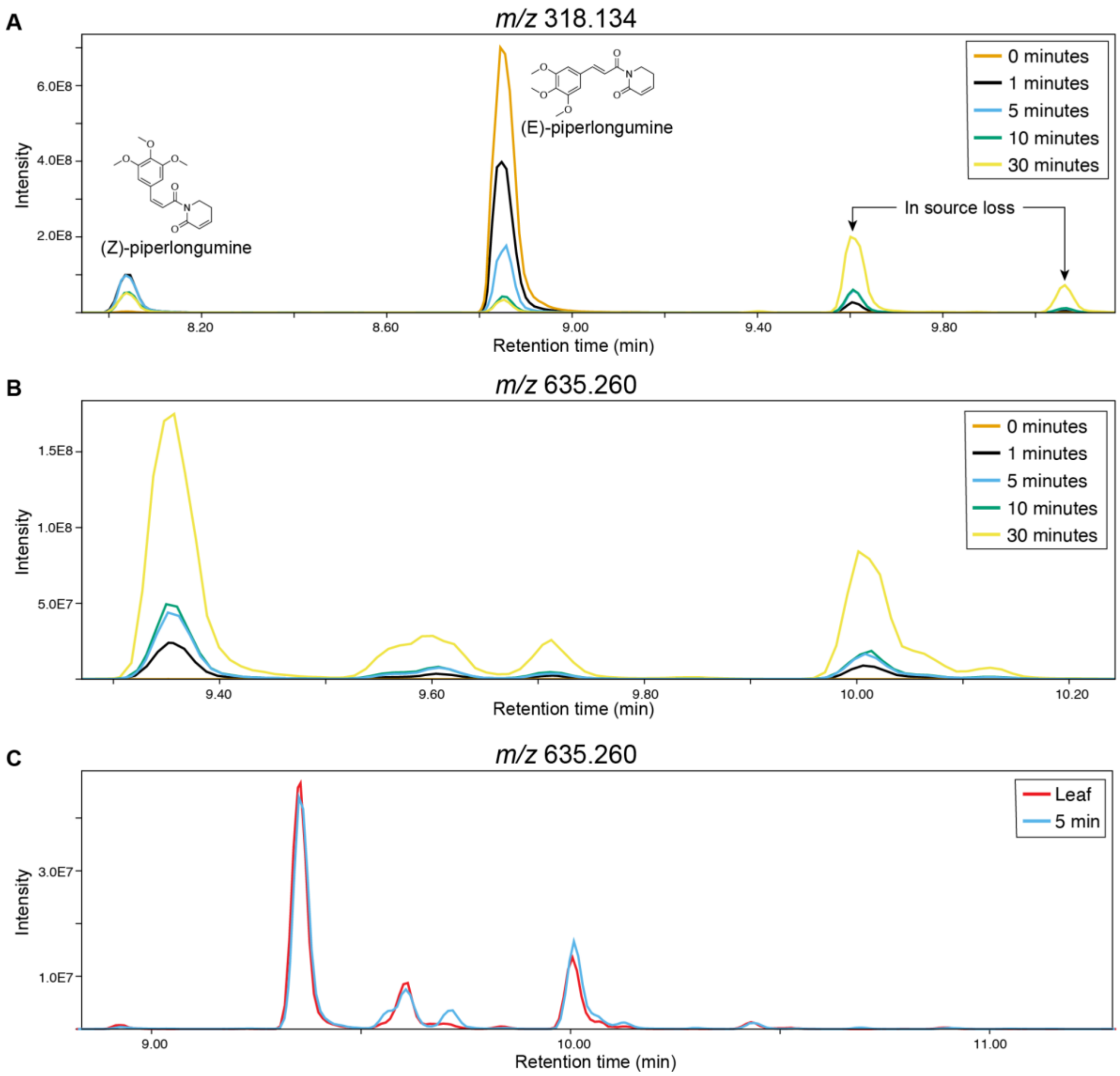


**S. Figure 7**. Results of the direct LC-MS analysis of the piperlongumine dimer reaction mixture (panel A and B). Extracted ion chromatogram of *m/z* 318.134 (i.e., $[M+H]^+$ adduct of piperlongumine monomer) and *m/z* 635.260 (i.e., $[M+H]^+$ adduct of piperlongumine dimer) are shown in panel A and B, respectively. The different time points of the reaction (see **Experimental section**) are displayed with different colours. Panel C shows an overlay of the extracted ion chromatogram of *m/z* 635.260 in the leaf organ and reaction mixture (5 minutes time point).

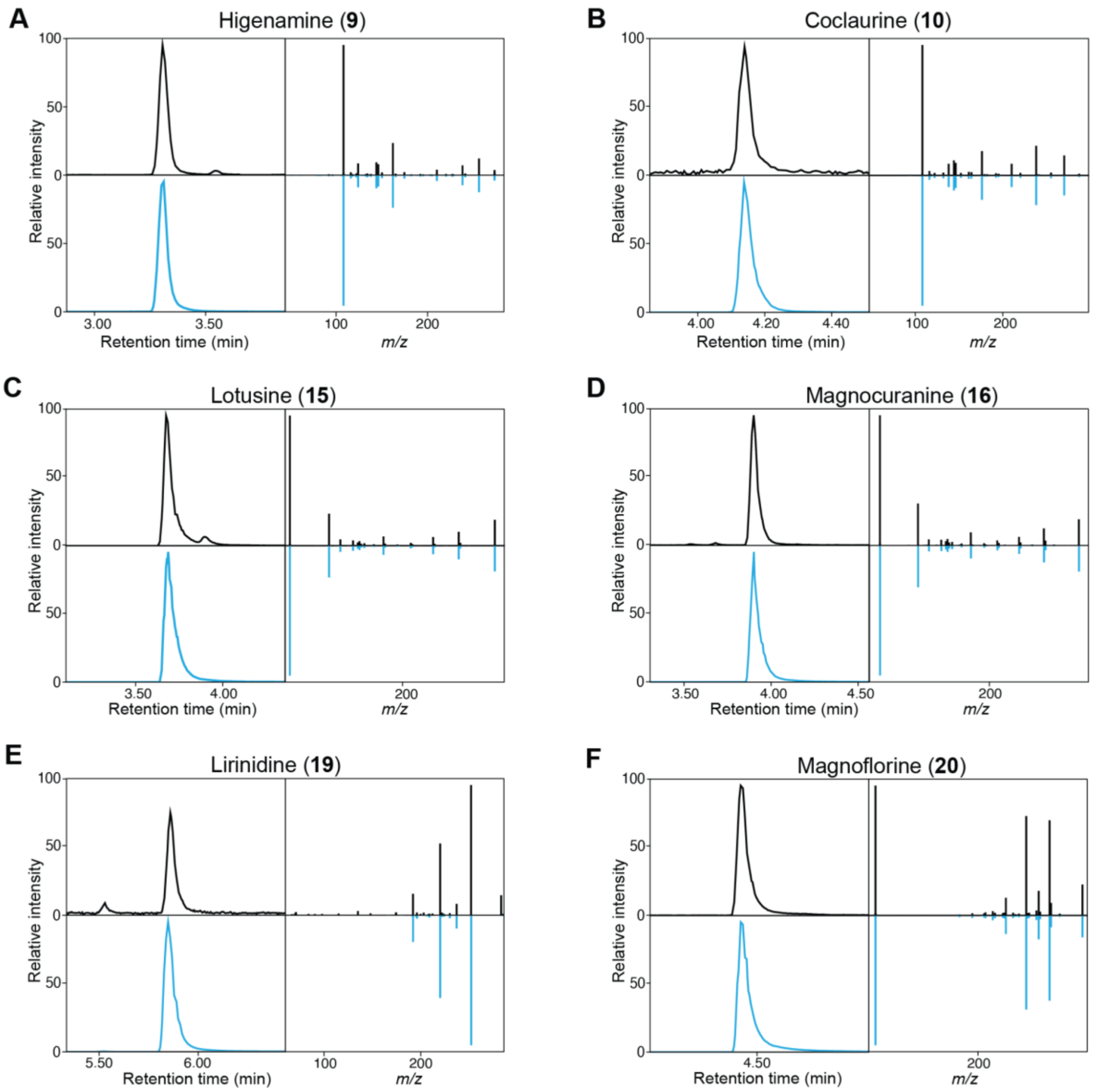


**S. Figure 8**. Confirmation of BIAs and aporphine alkaloids via retention time matching with analytical standards. Extracted ion chromatograms and MS/MS spectra of sample (top) and standards (bottom) are coloured in black and blue, respectively. A) Higenamine (**9**), *m/z* 272.129; B) Coclaurine (**10**), *m/z* 286.143; C) Lotusine (**15**), *m/z* 314.175; D) Magnocurarine (**16**), *m/z* 314.175; E) Lirinidine (**17**), *m/z* 282.149; F) Magnoflorine (**18**), m/z 342.171.

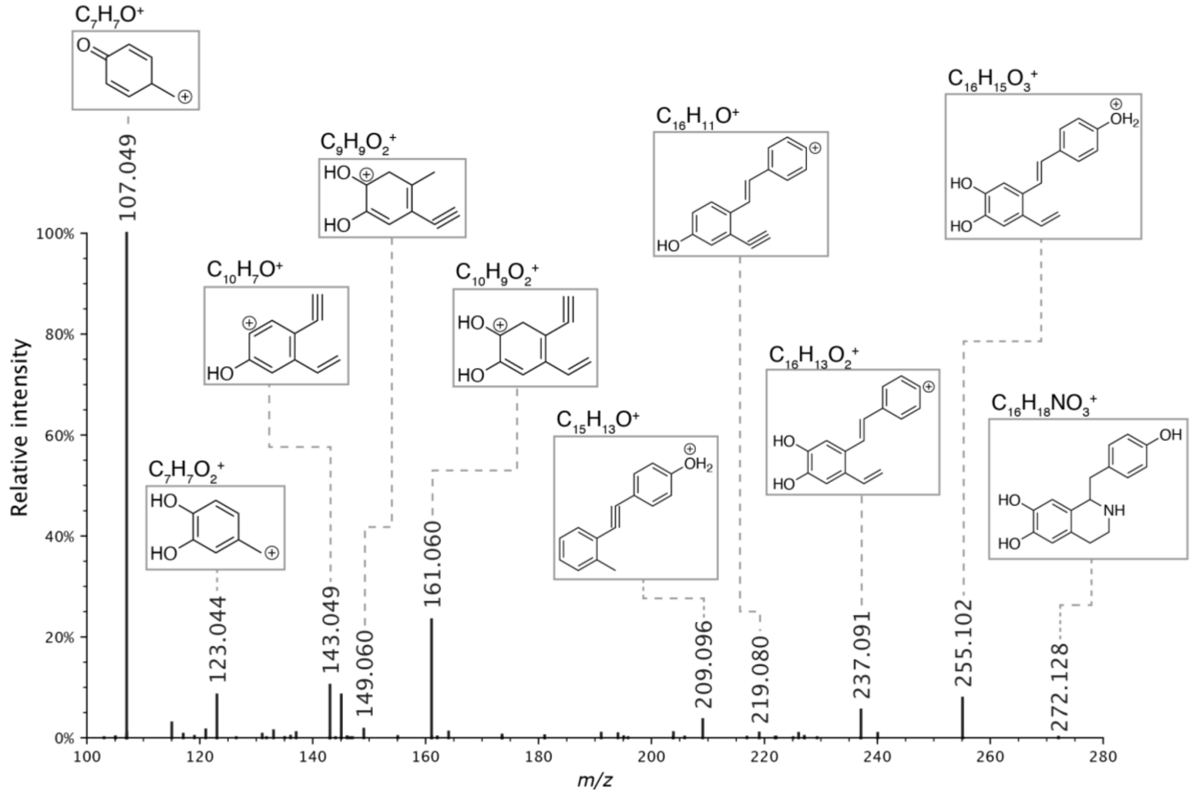


**S. Figure 9**. Proposed MS/MS fragmentation pathway for higenamine (in accordance with (Zhao et al. 2022)). The MS/MS spectrum in the figure can be reproduced using the Metabolomics Spectrum Resolver at this link.

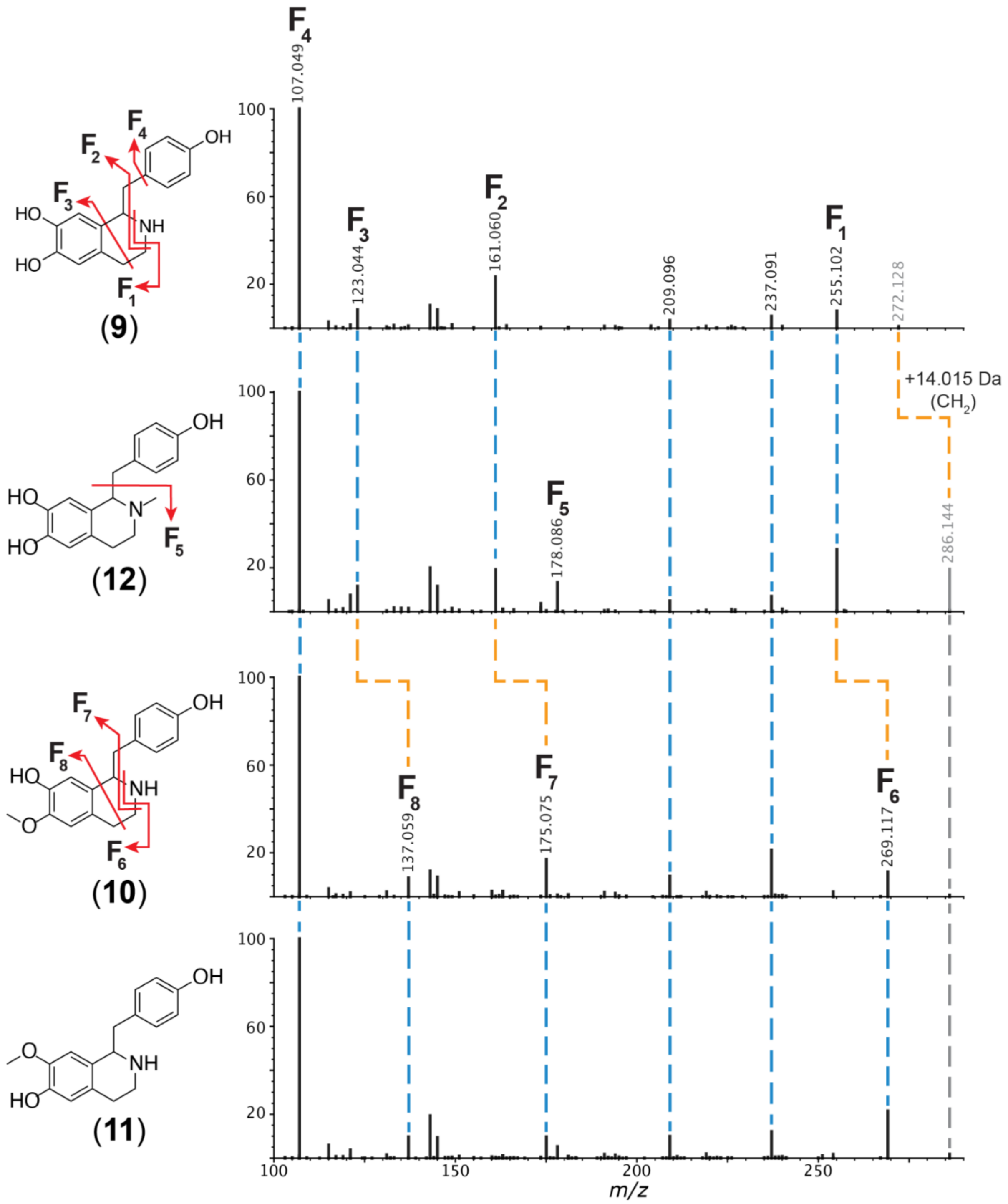


**S. Figure 10**. Interpretation of BIAs MS/MS spectra. Dashed gray lines represent precursor *m/z*. Dashed blue lines represent matching fragment peaks. Dashed orange lines represent matching fragment peaks shifted by the same difference as the precursors' *m/z* (+14.015 Da). Proposed fragmentation pathways are represented by red arrows in the chemical structures. MS/MS spectra in the figure can be reproduced using the Metabolomics Spectrum Resolver at the following links: higenamine, N-methylhigenamine, coclaurine, isococlaurine.

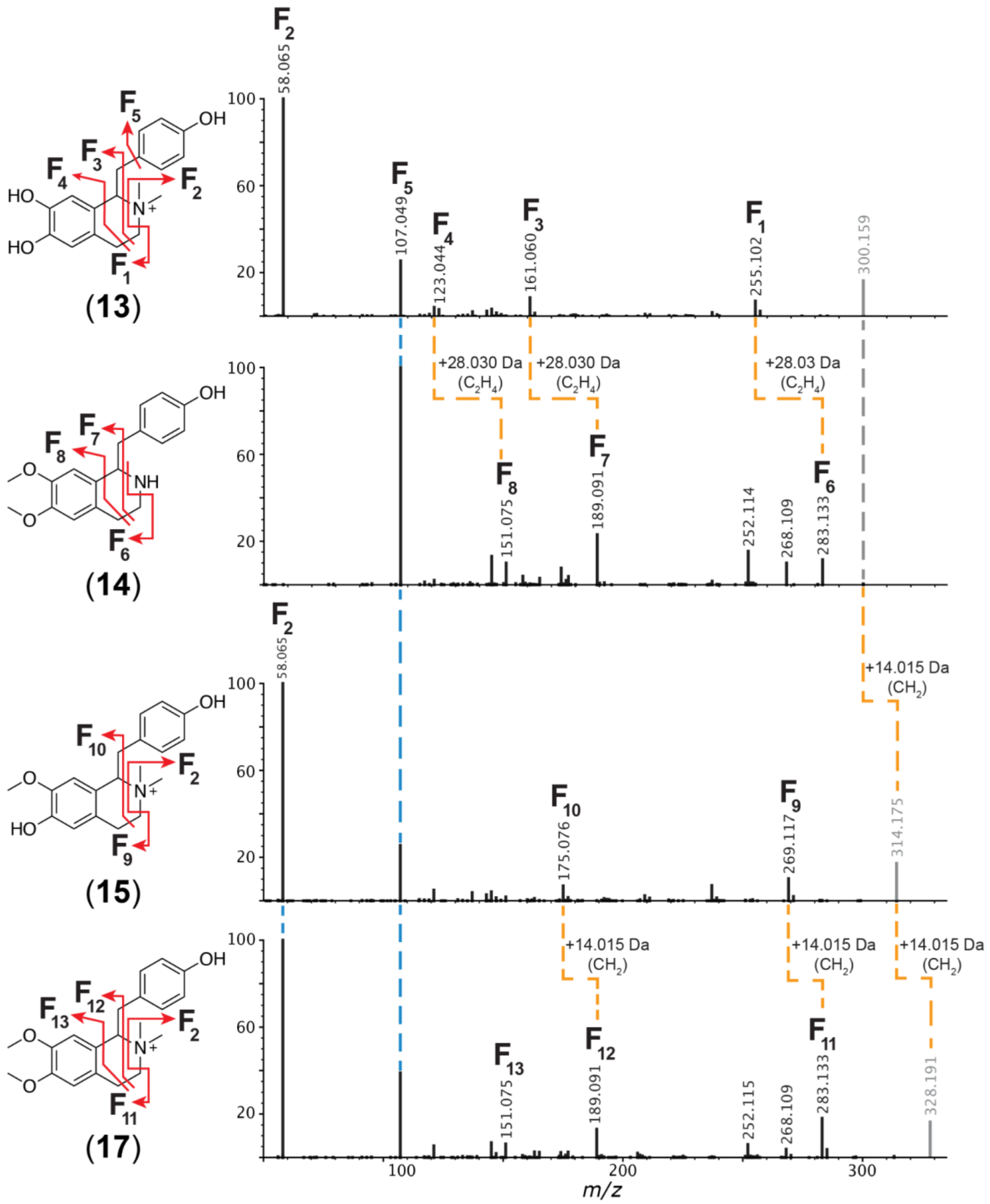


**S. Figure 11**. Interpretation of BIAs MS/MS spectra. Dashed gray lines represent precursor *m/z*. Dashed blue lines represent matching fragment peaks. Dashed orange lines represent matching fragment peaks shifted by the same difference as the precursors' *m/z* (+14.015 Da and +28.030). Proposed fragmentation pathways are represented by red arrows in the chemical structures. MS/MS spectra in the figure can be reproduced using the Metabolomics Spectrum Resolver at the following links: N-dimethylhigenamine, Norarmepavine, Lotusine, N-methylarmepavine

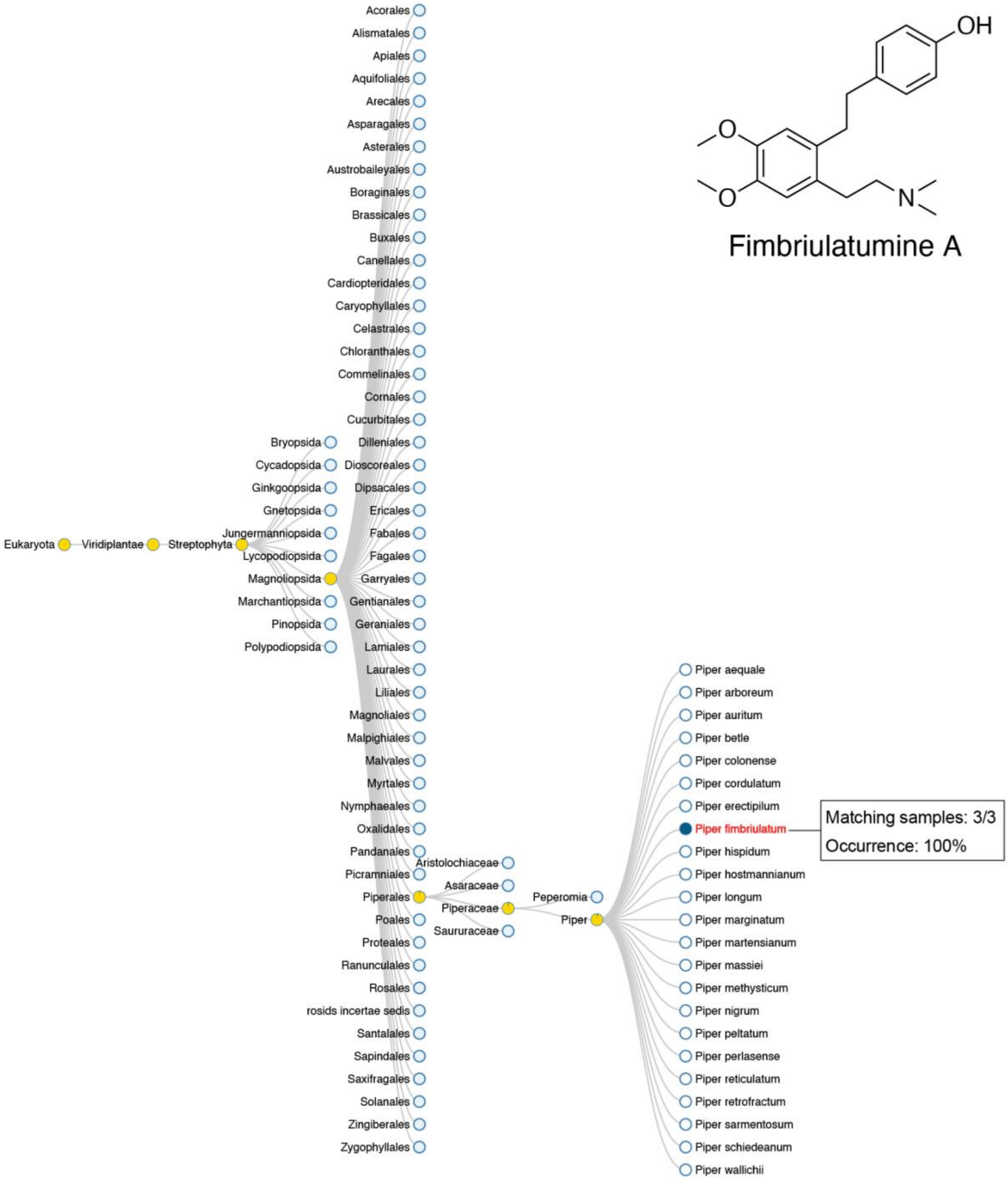


**S. Figure 12**. plantMASST search output of cuspidatin (**22**). For each taxonomic level, pie charts display the proportion of samples in plantMASST database for which an MS/MS match was retrieved (blue: matches; yellow: no matches). MS/MS matches were only retrieved in all (*n*=3) LC-MC/MS data linked to *P. fimbriulatum* in the plantMASST reference database. PlantMASST search results can be reproduced here.

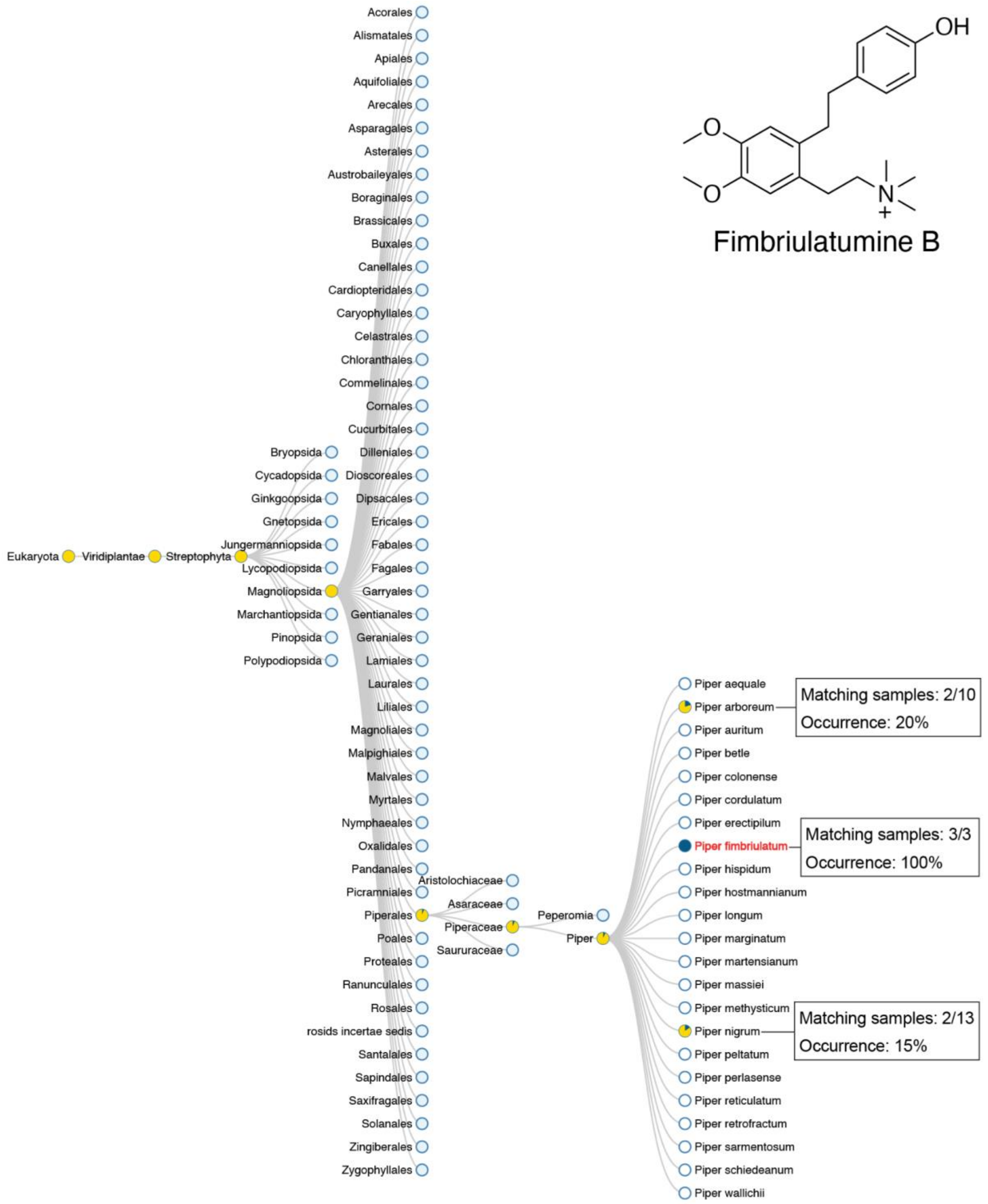


**S. Figure 13**. plantMASST search output of fimbriulatumine (**23**). For each taxonomic level, pie charts display the proportion of samples in plantMASST database for which an MS/MS match was retrieved (blue: matches; yellow: no matches). MS/MS matches were only retrieved in all ($n$=3) LC-MC/MS datafiles linked to *P. fimbriulatum* in the plantMASST reference database. MS/MS matches were also retrieved in 2 LC-MC/MS datafiles linked to *P. arboreum* and *P. nigrum*. Such datafiles belong to a MassIVE dataset uploaded by our own lab (MSV000087844) and further inspection of the raw data revealed that carry-over occurred during the analysis, (probably due to the positive charge carried by the molecule which causes strong peak tailing). This caused the detection of fimbriulatumine B in these two species, although not naturally present. PlantMASST search results can be reproduced here.

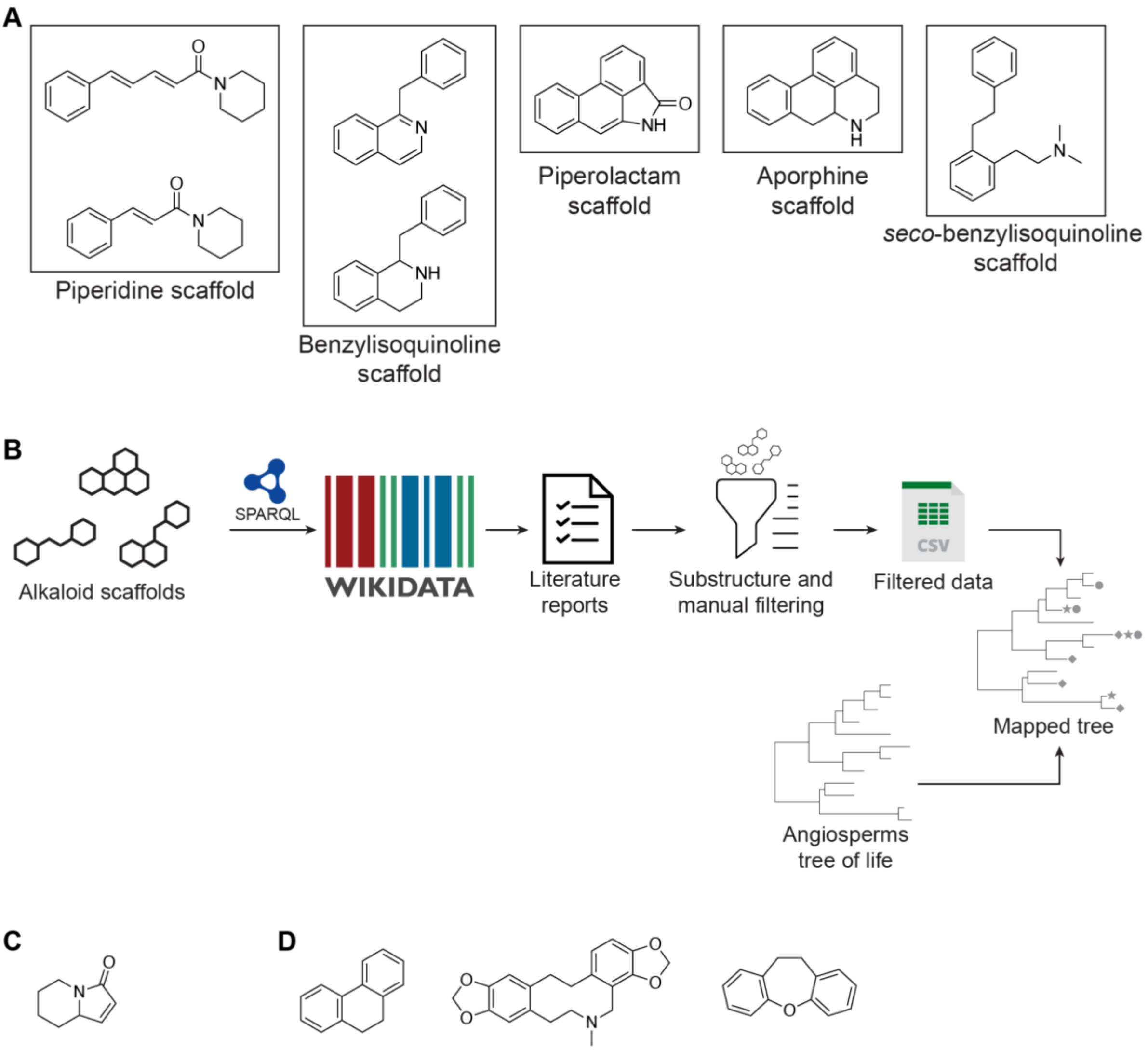


**S. Figure 14**. Computational workflow used to map literature reports for the alkaloid scaffolds found in *P. frimbriulatum* in the present study onto the Angiosperm tree of life(Zuntini et al. 2024). A) Chemical structures used to search Wikidata for literature reports; B) Schematic representation of the workflow; C) Chemical (sub)structures removed from the Wikidata query results for the piperidine scaffold; D) Chemical (sub)structures removed from Wikidata query results for the *seco*-benzylisoquinoline scaffold

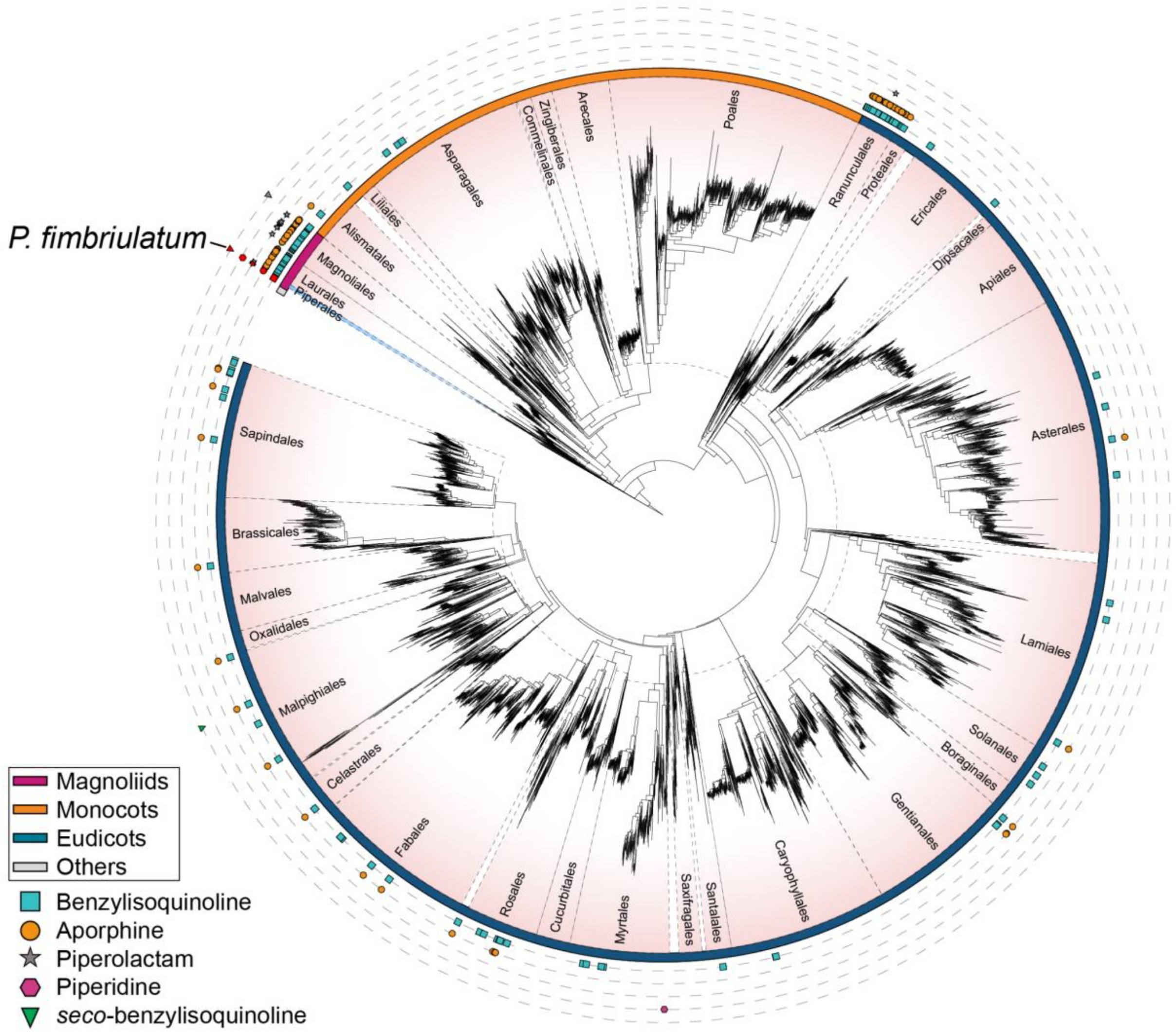


**S. Figure 15**. Angiosperm tree of life from (Zuntini et al. 2024) mapped with literature reports, mined from Wikidata, of the alkaloid scaffolds considered in the present study (i.e., benzylisoquinoline, aporphine, piperolactam, piperidine, *seco*-benzylisoquinoline). Each leaf in the tree corresponds to a representative species for each genus as described in the original publication. Different plant orders are separated by dashed black lines and the Piperales order is highlighted in light blue. Reports for each scaffold are represented with different colored shapes (see legend in the figure). Reports for the *Piper* genus are highlighted in red. Coloured arcs around the tree indicate the four main clades of Angiosperms as described in the original publication: Magnoliids, Monocots, Eudicots and ANA grade (i.e., Amborellales, Nymphaeales and Austrobaileyales). More details about the construction of the tree are provided in the **Experimental section**. The figure was created using iTOL(Letunic and Bork 2024). The original tree can be accessed at https://itol.embl.de/tree/14723112167277531728383616

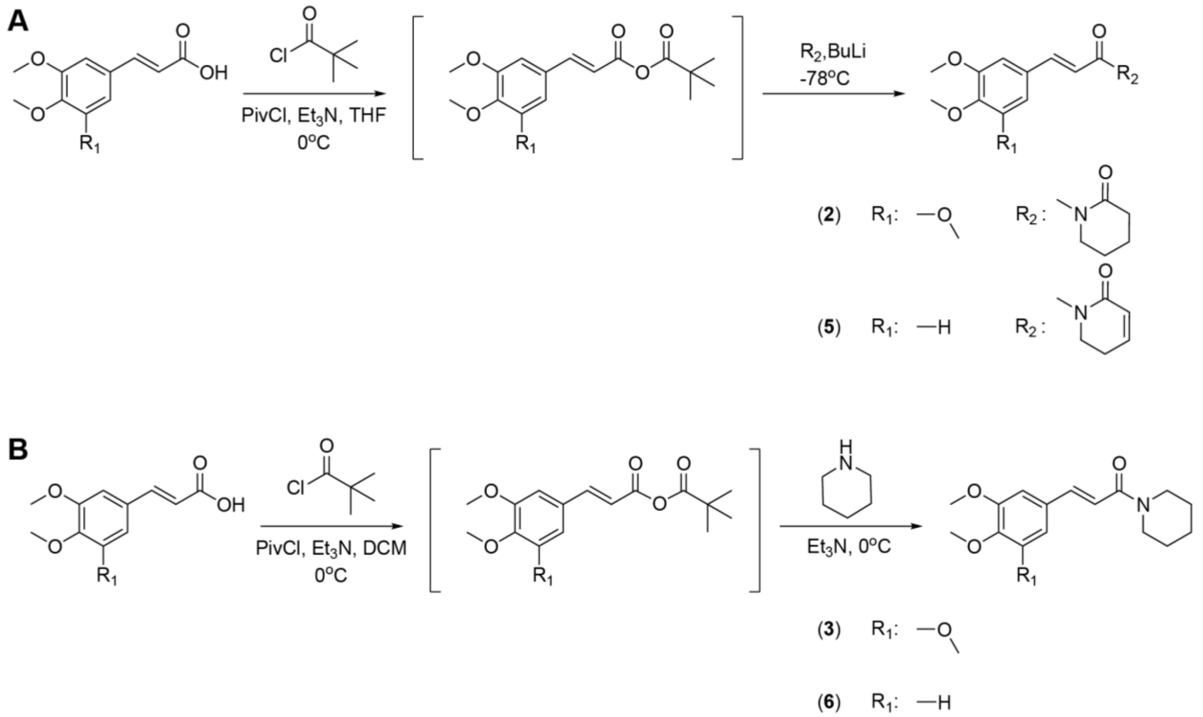


**S. Figure 16**. Scheme of the chemical synthesis of piperamides. A) For compounds (**2**) and (**5**), 3,4,5-Trimethoxy and 3,4-Dimethoxy cinnamic acid, respectively, were dissolved in dry tetrahydrofuran (THF) under argon atmosphere, added with triethylamine ($Et_3N$) and pivaloyl chloride (PivCl), and the mixture was stirred for 1 hour at 0 °C. Afterwards, 2-piperidinone ($R_2$) and *n*-butyllithium (BuLi) were added and stirred for 1h at −78 °C. Finally, the reaction mixture was quenched, extracted and purified.d; B) For compounds (**3**) and (**6**), 3,4,5-Trimethoxy and 3,4-Dimethoxy cinnamic acid, respectively, were dissolved in dichloromethane (DCM), added with $Et_3N$ and PivCl, and the mixture was stirred for 1 hour at 0 °C. Afterwards, $Et_3N$ was added to a stirred solution of piperidine in dry DCM, added to the reaction mixture and stirred overnight at 0 °C. Finally, the reaction mixture was quenched, extracted and purified. A more detailed description of the chemical synthesis is provided in the **Experimental Section** section.

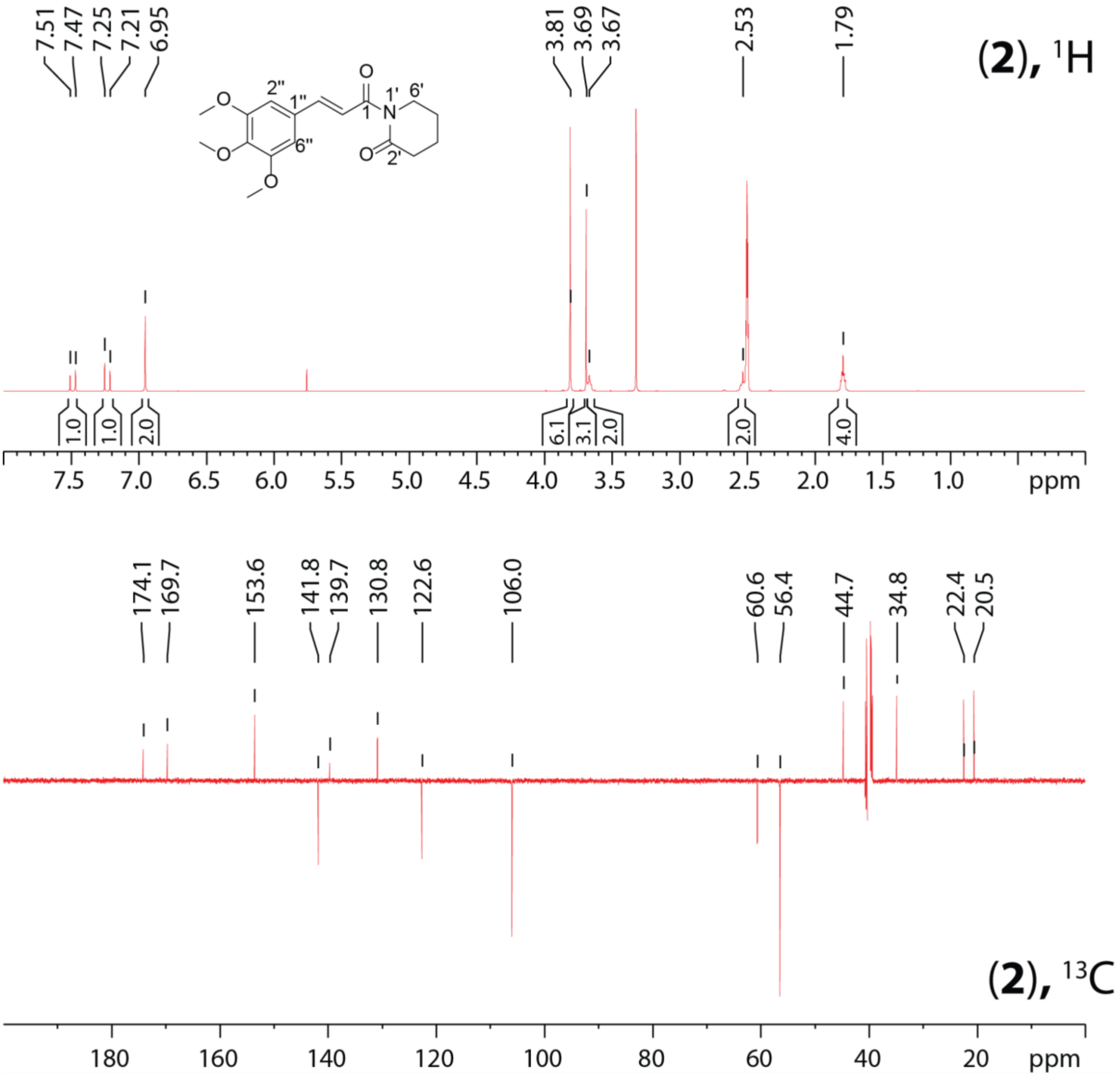


**S. Figure 17.** $^{1}H$ and $^{13}C$ NMR spectra of dihydropiperlongumine (**2**). $^{1}H$ NMR (top) (400.0 MHz, $C_2D_6OS$) δ = 7.49 (1H, d, 15.2 Hz , H-2), 7.23 (1H, d, 15.2 Hz,H-3), 6.95 (2H, s, H-2'', H-6''), 3.80 (6H, s, H-3''-OMe, H-5''-OMe), 3.69 (3H, s, H-4'' OMe), 3.66 (2H, m br, H-6'), 2.54 (2H, m br, H-3'), 1.80 (4H, q, 4 Hz, H-4', H-5'). The $^{1}H$ signal at 2.50 is $C_2H_6OS$; the $^{1}H$ signal at 3.33 is $H_2O$; the $^{1}H$ signal at 5.70 is $CH_2Cl_2$. $^{13}C$ NMR (bottom) (101 MHz, $C_2D_6OS$) δ = 174.1 (C, C-2'), 169.7 (C, C-1), 153.5 (C, C-3'', C-5''), 141.8 (CH, C-3), 139.7 (C, C-4''), 130.8 (C, C-1''), 122.6 (CH, C-2), 106.0 (CH, C-2'', C-6''), 60.6 (CH3, C-4''OMe), 56.4 (CH3, C3''OMe, C5''OMe),44.7 (CH2, C-6'), 34.8 (CH2, C-3'), 22.4 (CH2, C-5'), 20.5 (CH2, C-4'). The $^{13}C$ signal at 39.52 is $C_2D_6OS$.

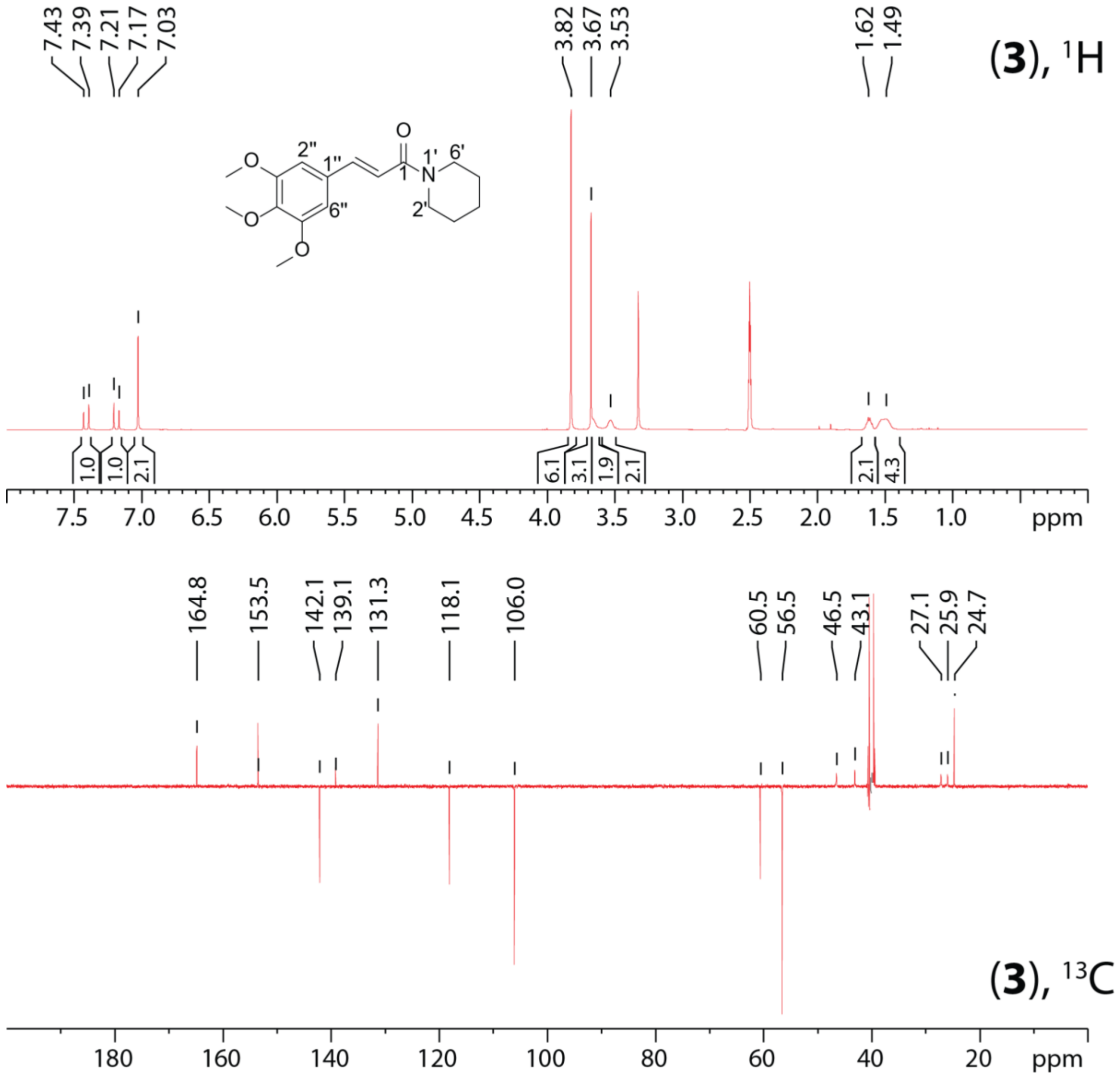


**S. Figure 18**. $^{1}$H and $^{13}$C NMR spectra of  of 1-(3,4,5-Trimethoxycinnamoyl)piperidine (**3**). $^{1}$H NMR (top) (400.0 MHz, $C_2D_6OS$) δ = 7.41 (1H, d, 15.2 Hz , H-2), 7.19 (1H, d, 15.2 Hz,H-3), 7.03 (2H, s, H-2'', H-6''), 3.82 (6H, s, H-3''-OMe, H-5''-OMe), 3.68 (3H, s, H-4''  OMe), 3.65 (2H, m br, H-6'), 3.53 (2H, m br, H-2'), 1.62 (2H, q, 4 Hz, H-4'), 1.50 (4H, m br, H-3', H-5'). The $^{1}$H signal at 2.50 is $C_2H_6OS$; the $^{1}$H signal at 3.33 is $H_2O$; the $^{1}$H signal at 5.70 is $CH_2Cl_2$. $^{13}$C NMR (bottom) (101 MHz, $C_2D_6OS$) δ =  164.8 (C, C-1), 153.5 (C, C-3'', C-5''), 142.1 (CH, C-3), 139.1 (C, C-4''), 131.3 (C, C-1''), 118.1 (CH, C-2), 106.0 (CH, C-2'', C-6''), 60.5 (CH3, C-4''OMe), 56.5 (CH3, C3''OMe, C5''OMe), 46.5 (CH2, C-2'), 43.1 (CH2, C-6'), 27.1 (CH2, C-3'), 25.9 (CH2, C-5'), 24.7 (CH2, C-4'). The $^{13}$C signal at 39.52 is $C_2D_6OS$.

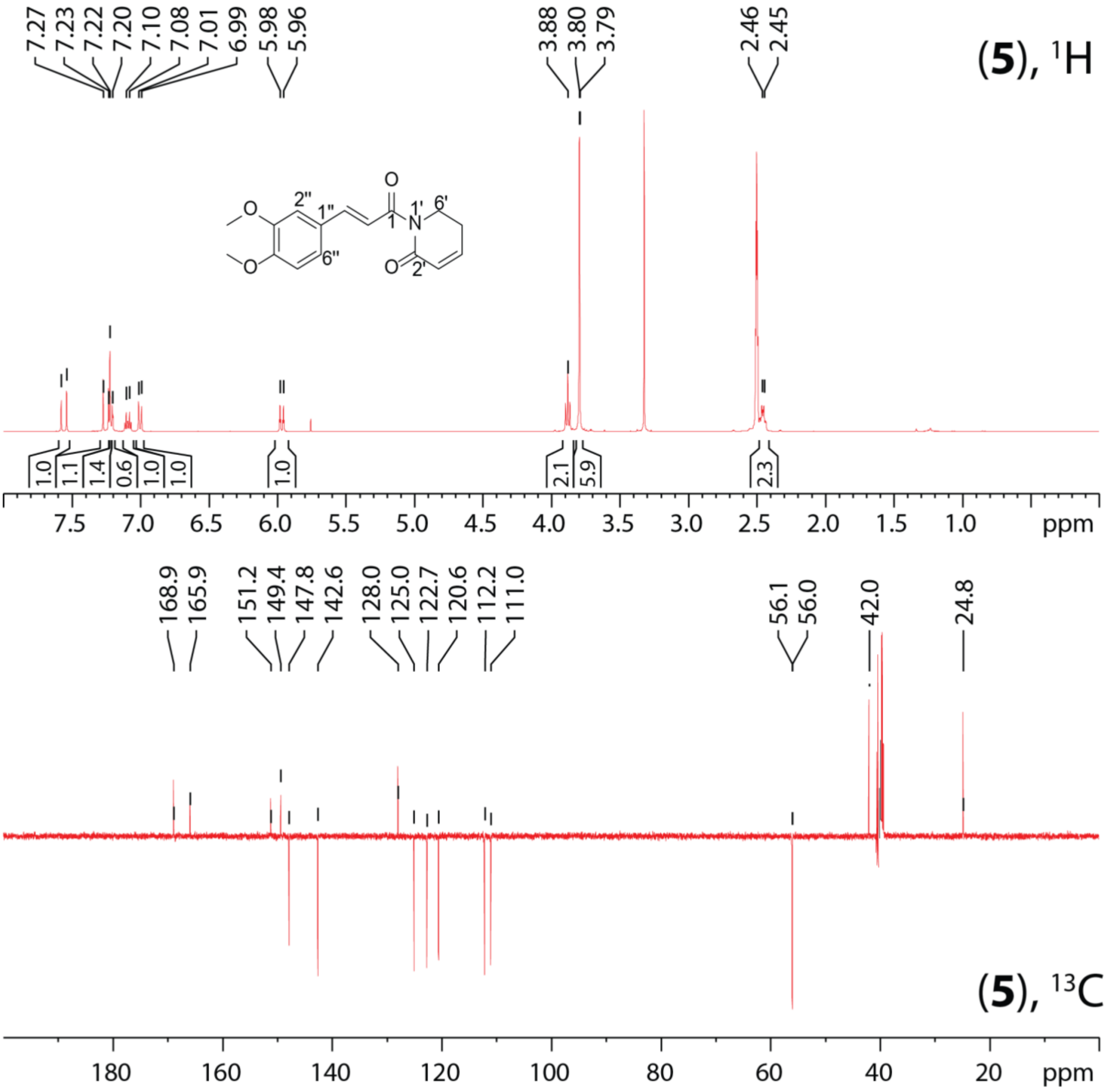


**S. Figure 19**. $^{1}H$ and $^{13}C$ NMR spectra of 3'-Demethoxypiperlongumine (**5**). $^{1}H$ NMR (top) (400.0 MHz, $C_2D_6OS$) δ = 7.56 (1H, d, 15.2 Hz , H-2), 7.25 (1H, d, 15.2 ,H-3), 7.22 (1H, d, 2.0 Hz ,H-2''), 7.21 (1H, dd, 2, 8.4 Hz, H-6''), 7.09 (1H, td, 4, 8.6 Hz, H-4'), 7.00 (1H, d, 8.2 Hz, H-5''), 5.97 (1H, td, 1.6, 9.7 Hz, H-3'), 3.88 (2H, t, 6.5 Hz, H-6'), 3.80 (3H, s, H-3''OMe), 3.79 (3H, s, H-4''OMe), 2.46 (2H, m br, H-5'). The $^{1}H$ signal at 2.50 is $C_2H_6OS$; the $^{1}H$ signal at 3.33 is $H_2O$; the $^{1}H$ signal at 5.70 is $CH_2Cl_2$. $^{13}C$ NMR (bottom) (101MHz, $C_2D_6OS$) δ = 168.9 (C, C-1), 165.9 (C, C-2'), 151.2 (C, C-3''), 149.4 (C, C-4''), 147.9 (CH, C-3'), 142.6 (CH, C-3), 128.0 (C, C-1''), 125.0 (C, C-4'), 122.7 (CH, C-6''), 120.5 (CH, C-2), 112.2 (CH, C-5''), 111.0 (C, C-2''), 56.0 (CH3, C-3''OMe), 56.0 (CH3, C4''OMe), 42.0 (CH2, C-6'), 24.8 (CH2, C-5'). The $^{13}C$ signal at 39.52 is $C_2D_6OS$.

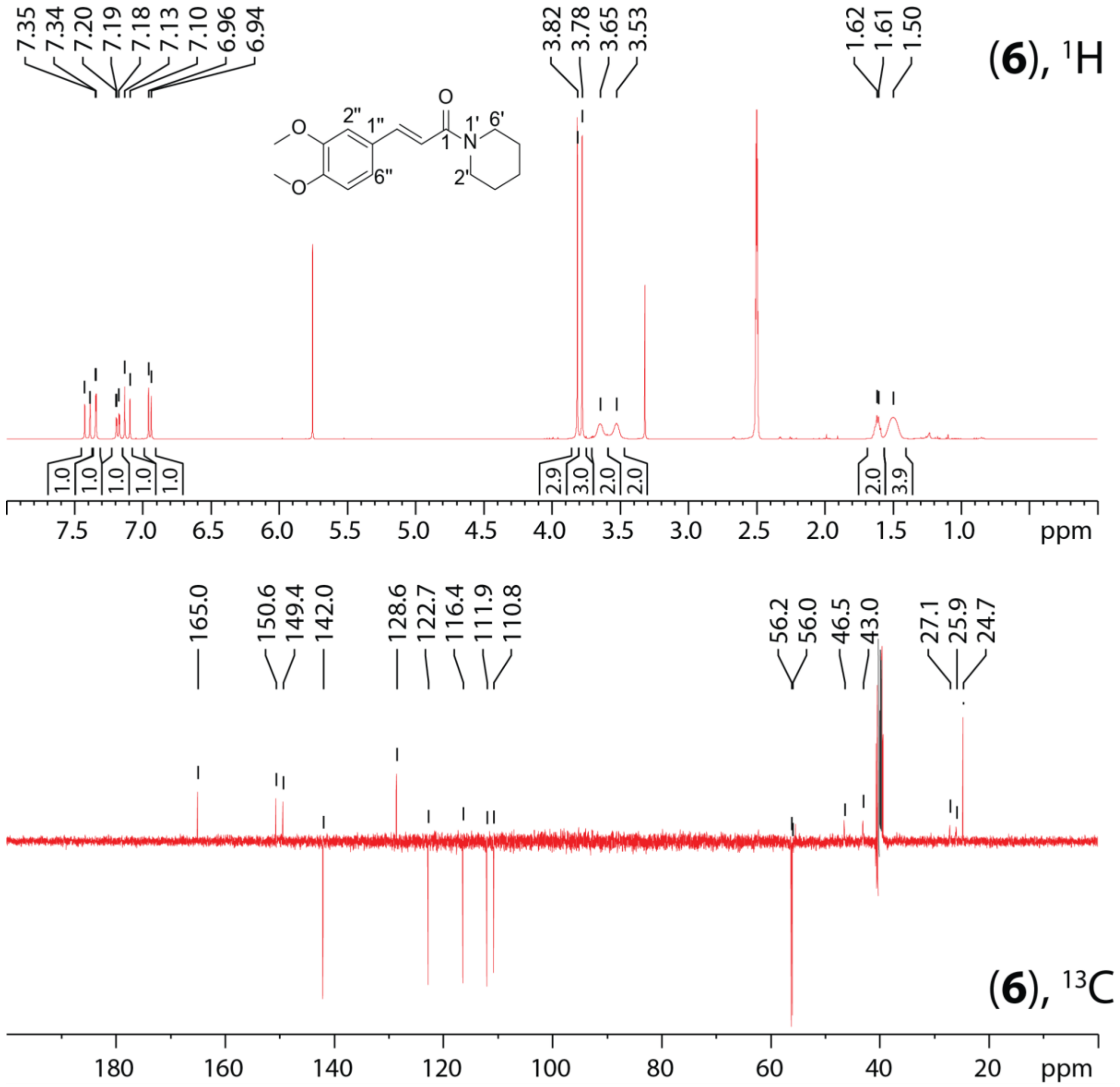


**S. Figure 20**. $^{1}$H and $^{13}$C NMR spectra of 1-(3,4-Dimethoxycinnamoyl)piperidine (**6**). $^{1}$H NMR (top) (400.0 MHz, $C_2D_6OS$) δ = 7.40 (1H, d, 15.2 Hz , H-2), 7.35 (1H, d, 2.0 Hz ,H-2''), 7.18 (1H, dd, 2, 8.4 Hz, H-6''), 7.11 (1H, d, 15.2 Hz, H-3), 6.95 (1H, d, 8.2 Hz, H-5''), 3.82 (3H, s, H-3''OMe), 3.78 (3H, s, H-4''OMe), 3.65 (2H, m br, H-6'), 3.53 (2H, m br, H-2'), ), 1.65 (2H, q, 4 Hz, H-4'), 1.50 (4H, m br, H-3', H-5'). The $^{1}$H signal at 2.50 is $C_2H_6OS$; the $^{1}$H signal at 3.33 is $H_2O$; the $^{1}$H signal at 5.70 is $CH_2Cl_2$. $^{13}$C NMR (bottom) (101 MHz, $C_2D_6OS$) δ = 165.0 (C, C-1), 150.6 (C, C-3''), 149.4 (C, C-4''), 142.0 (CH, C-3), 128.5 (C, C-1''), 122.6 (CH, C-6''), 116.4 (CH, C-2), 111.9 (CH, C-5''), 110.7 (CH, C-2''), 56.2 (CH3, C-3''OMe), 56.0 (CH3, C4''OMe), 46.5 (CH2, C-2'), 43.0 (CH2, C-6'), 27.1 (CH2, C-3'), 25.9 (CH2, C-5'), 24.7 (CH2, C-4'). The $^{13}$C signal at 39.52 is $C_2D_6OS$.

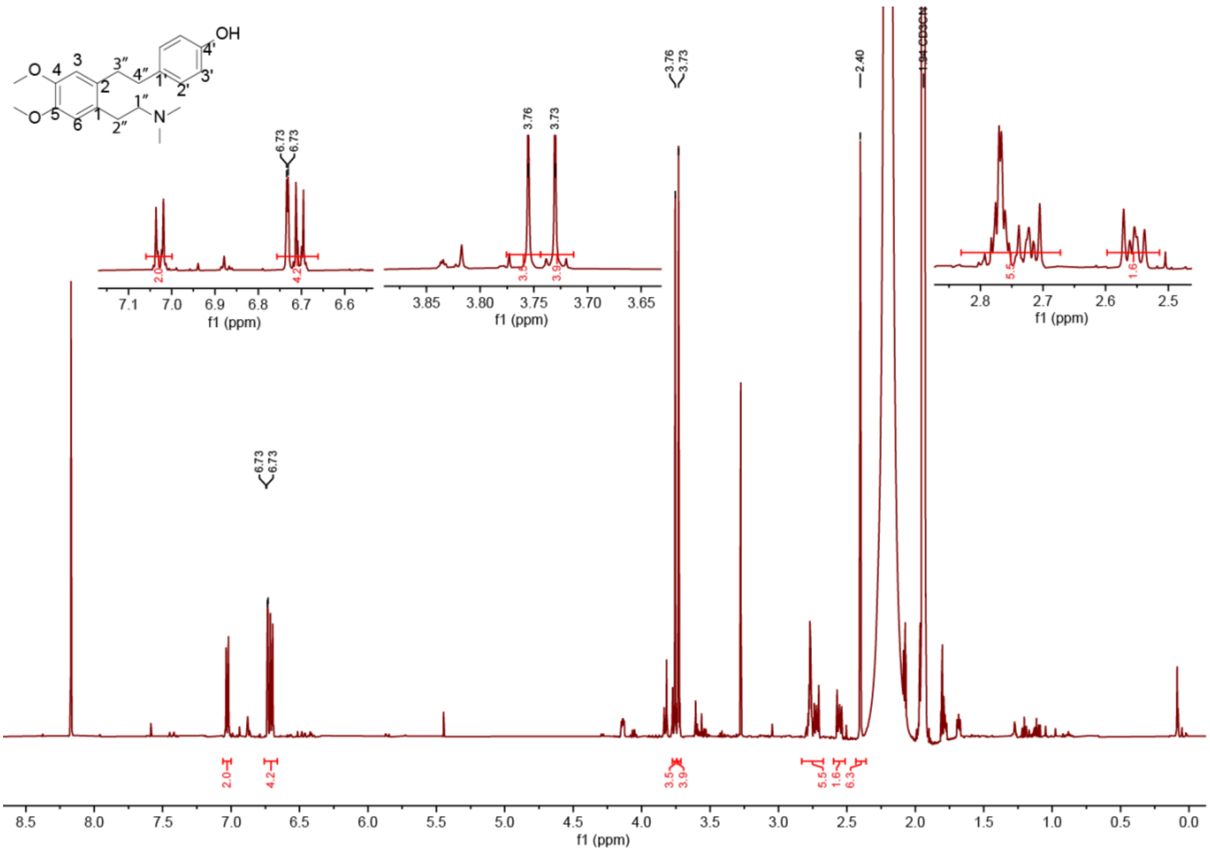


**S. Figure 21**. $^1$H NMR spectrum of cuspidatin (**22**). (500.0 MHz, $CD_3CN$) δ = 7.03 (m, 2H, H-2′), 6.73 (s, 1H, H-3), 6.73 (s, 1H, H6), 6.70 (m, 2H, H-3′), 3.76 (s, 3H, O–CH3), 3.73 (s, 3H, O–CH3), 2.70–2.79 (m, 6H, H-4’’, H-2’’, H-3’’), 2.56 (m, 2H,H-1’’), 2.40 (s, 6H, N–CH3). The sample contained small amounts of impurities which gave rise to signals around 1.68 and 1.78. The presence of such impurities did not interfere in structure annotation.

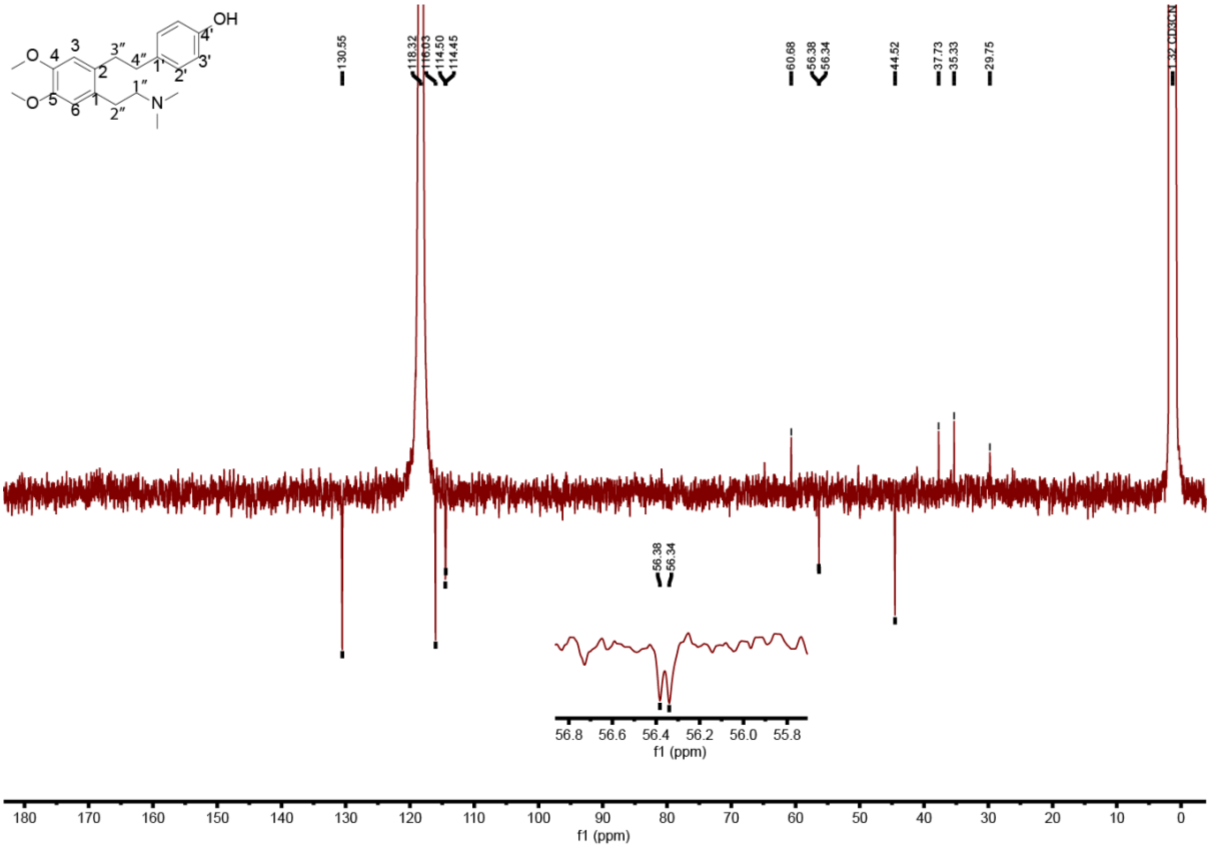


**S. Figure 22**. $^{13}$C NMR spectrum of cuspidatin (**22**) (125.7 MHz, CD3CN) δ = 156.1 (C-4′), 148.6 and 148.3 (C-4 and C-5), 134.0 (C-1′), 133.0 (C-2), 130.6 (C-2′ and C-6’), 130.0 (C-1), 116.0 (C-3′ and C-5’), 114.5 (C-3 or C-6), 60.7 (C-1’’), 56.4 (O–CH3), 56.3 (O–CH3), 44.5 (N–CH3), 37.7 (C-4’’), 35.3 (C-3’’), 29.8 (C-2’’). Not all signals were observable through 13C NMR due to the low amount of the compound. However, all carbons could be identified through HMBC.

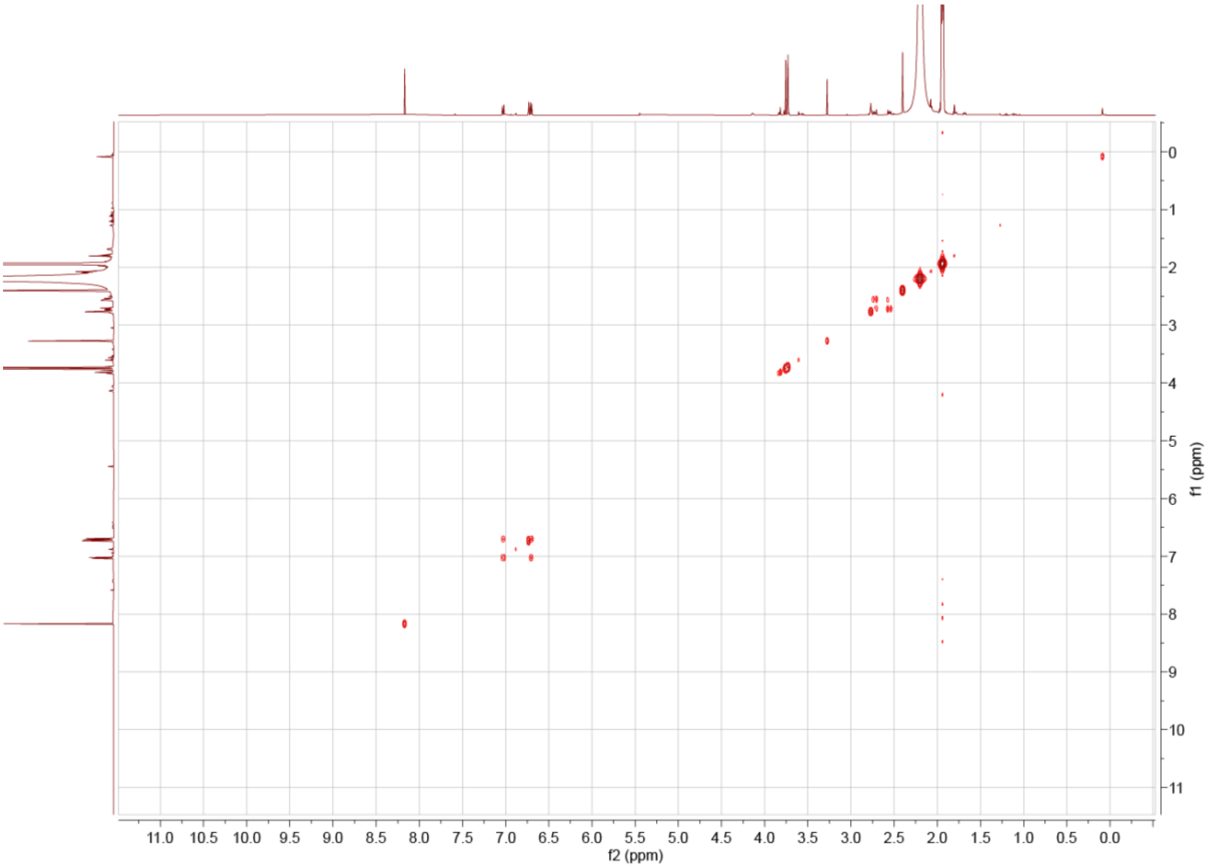


**S. Figure 23**. COSY spectrum of cuspidatin (**22**).

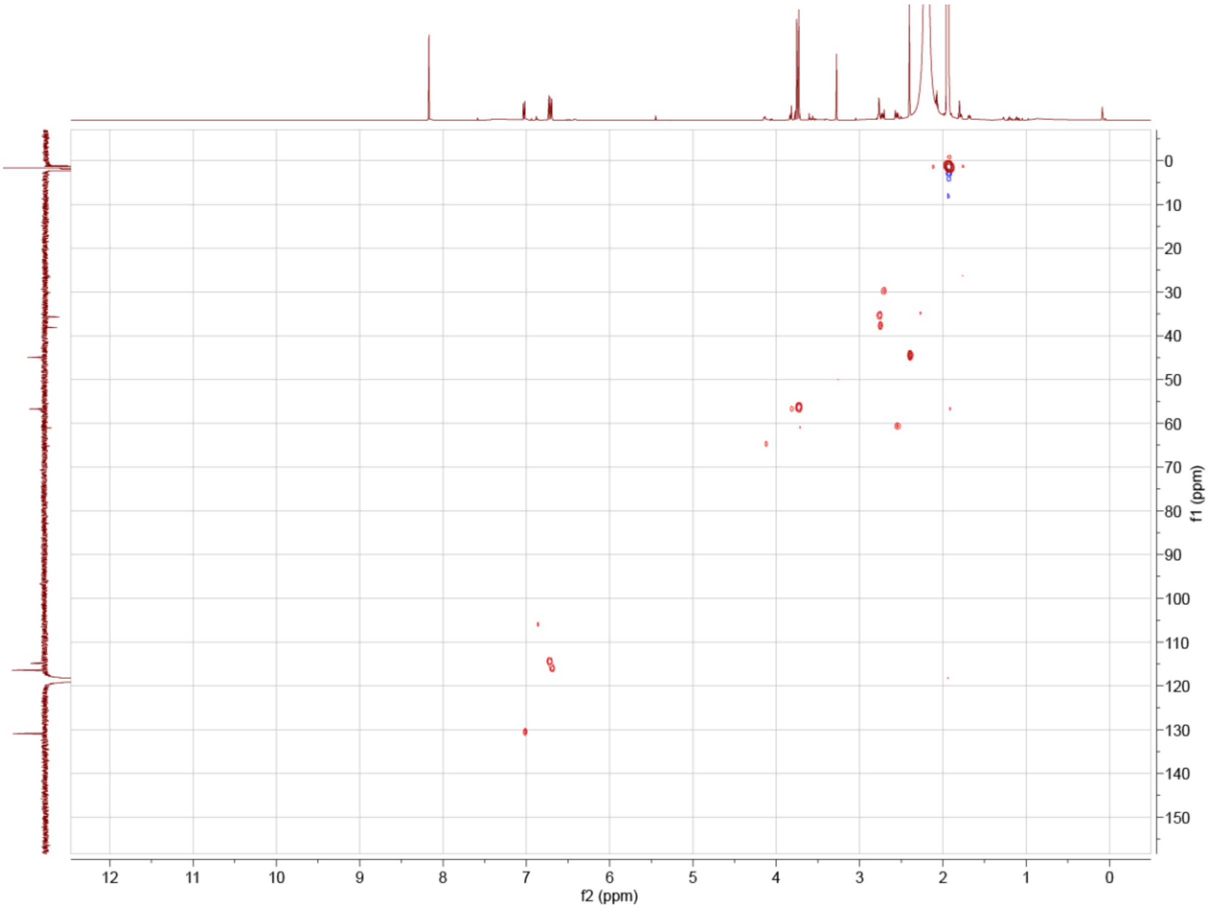

**S. Figure 24**. HSQC spectrum of cuspidatin (**22**).

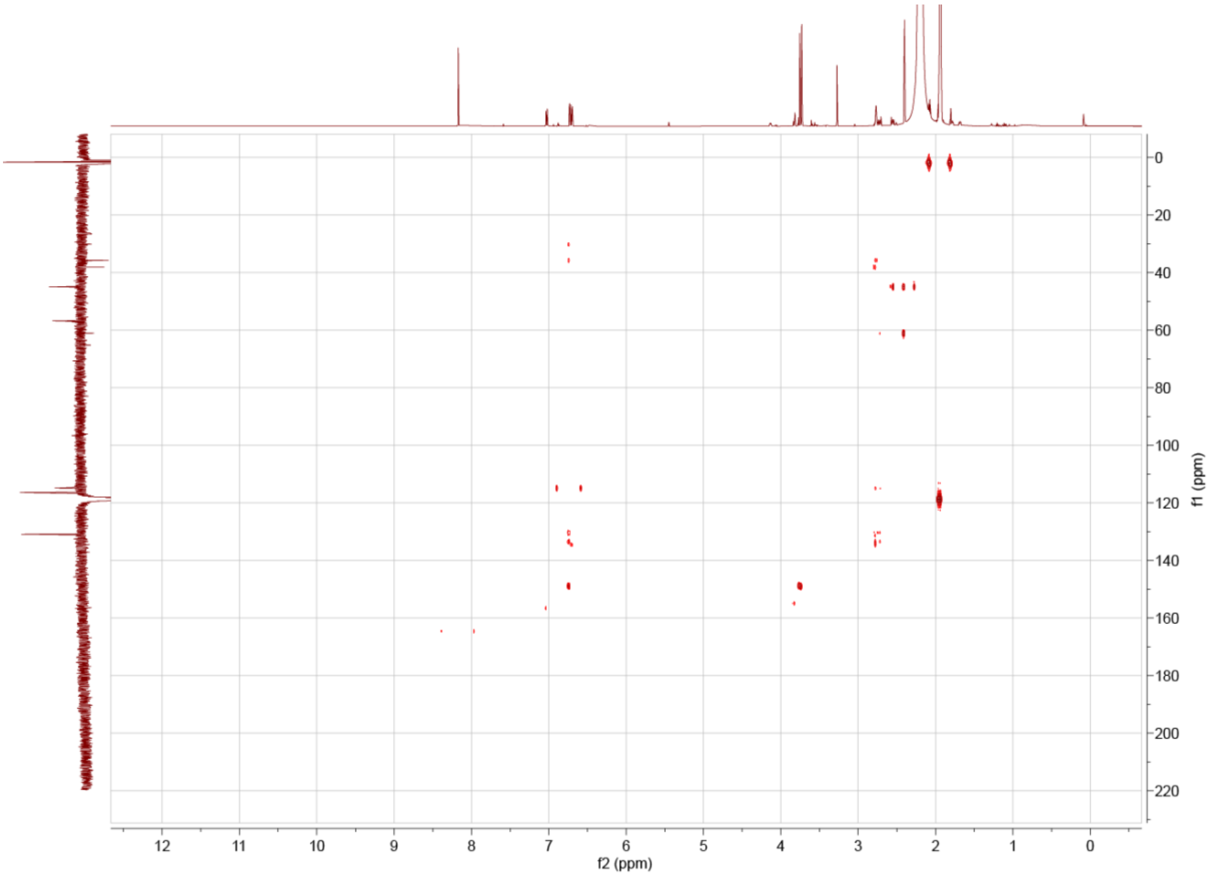


**S. Figure 25**. HMBC spectrum of cuspidatin (**22**).

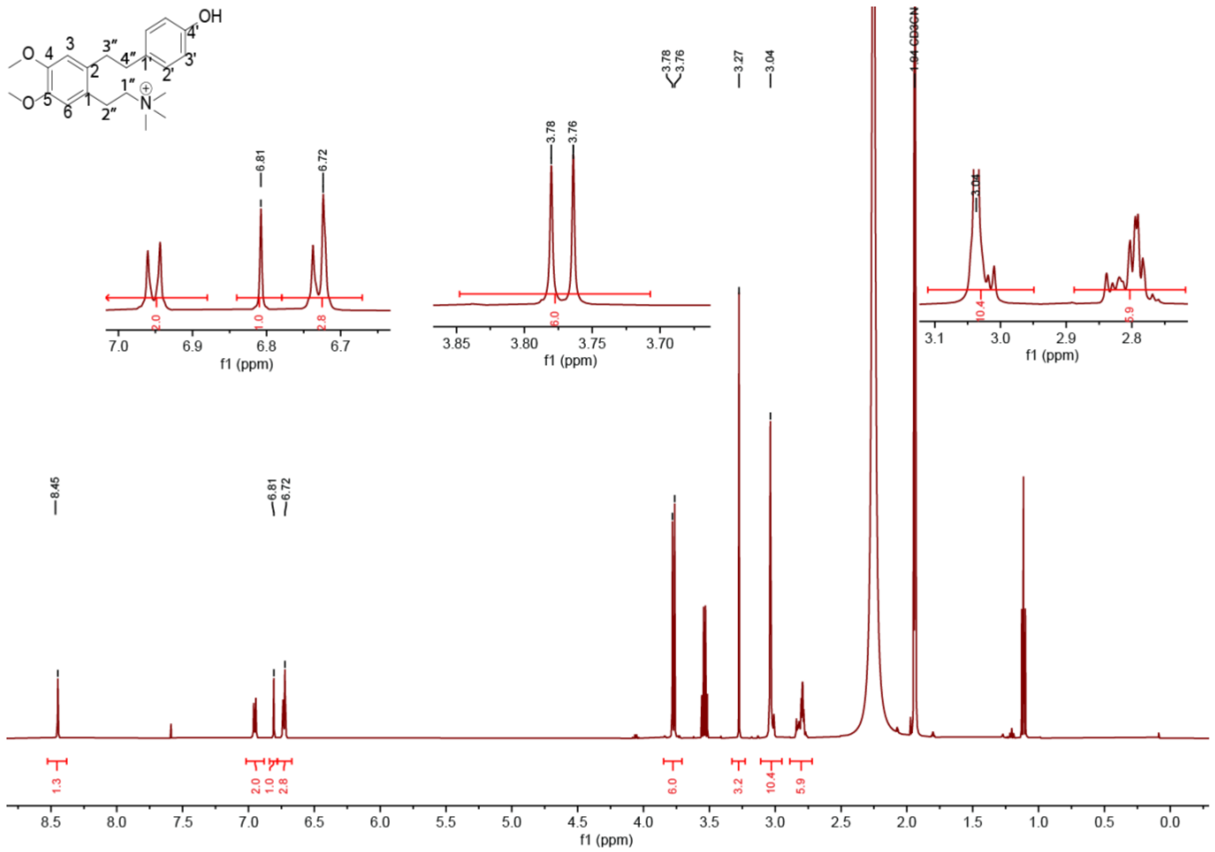


**S. Figure 26**. $^{1}$H NMR spectrum of fimbriulatumine (**23**). $^{1}$H NMR (top) (500.0 MHz, $CD_3CN$) δ = 6.95 (m, 2H, H2′), 6.81 (s, 1H, H-3), 6.73 (m, 2H, H-3′), 6.72 (s, 1H, H-6), 3.78 (s, 3H, O–CH3), 3.76 (s, 3H, O–CH3), 3.04 (s, 9H, N–CH3), 3.02 (m, 2H, H-1''), 2.76–2.84 (m, 6H, H-4'', H-2'', H-3''). Signals at 3.27 and 1.11 correspond to residual solvents methanol and ethanol, respectively.

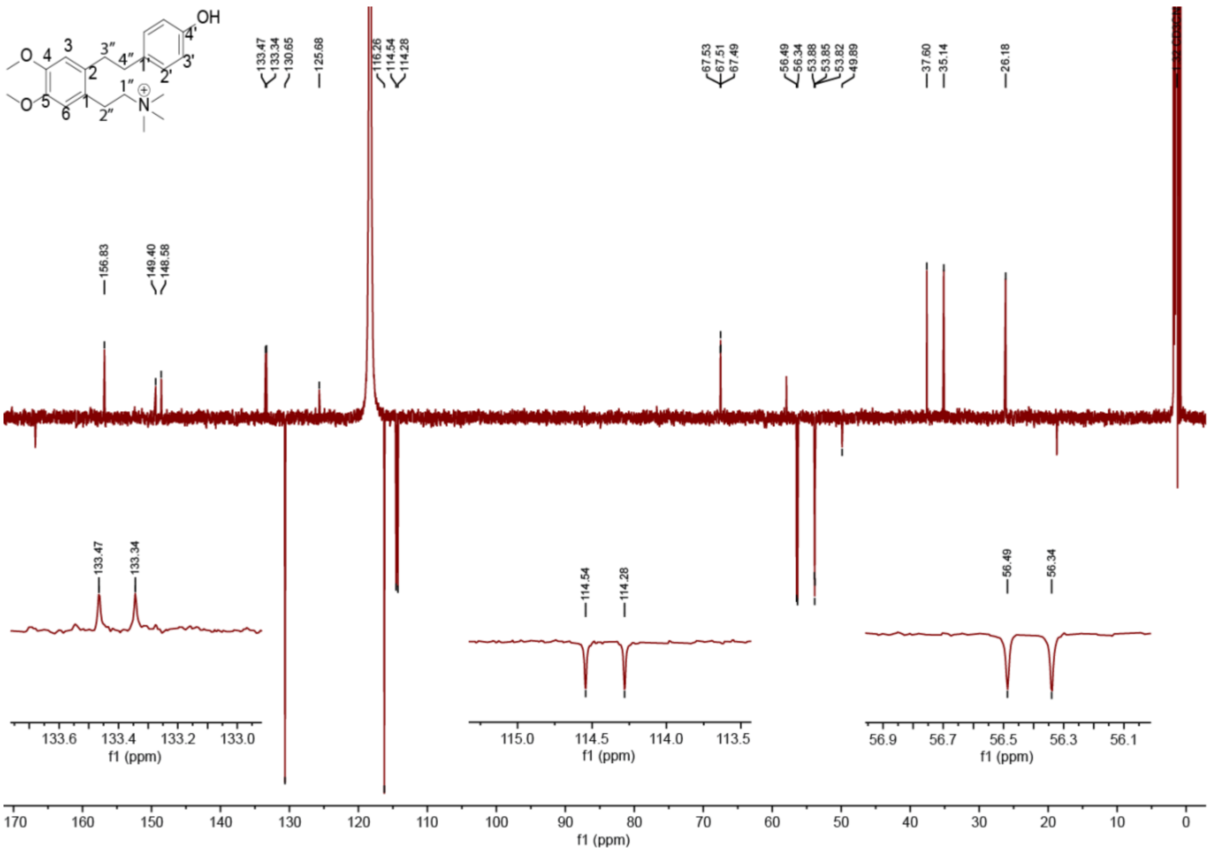


**S. Figure 27**. [13]C NMR spectrum of fimbriulatumine (**23**) (125.7 MHz, CD3CN) δ = 156.8 (C-4′), 149.4 and 148.6 (C-4 and C-5), 133.5 and 133.3 (C-2 and C-1′), 130.7 (C-2′ and C-6’), 125.7 (C-1), 116.3 (C-3′ and C-5’), 114.5 (C-3), 114.3 (C-6), 67.5 (m, C-1’’), 56.5 (O–CH3), 56.3 (O–CH3), 53.9 (m, N–CH3), 37.6 (C-4’’), 35.1 (C-3’’), 26.2 (C-2’’).

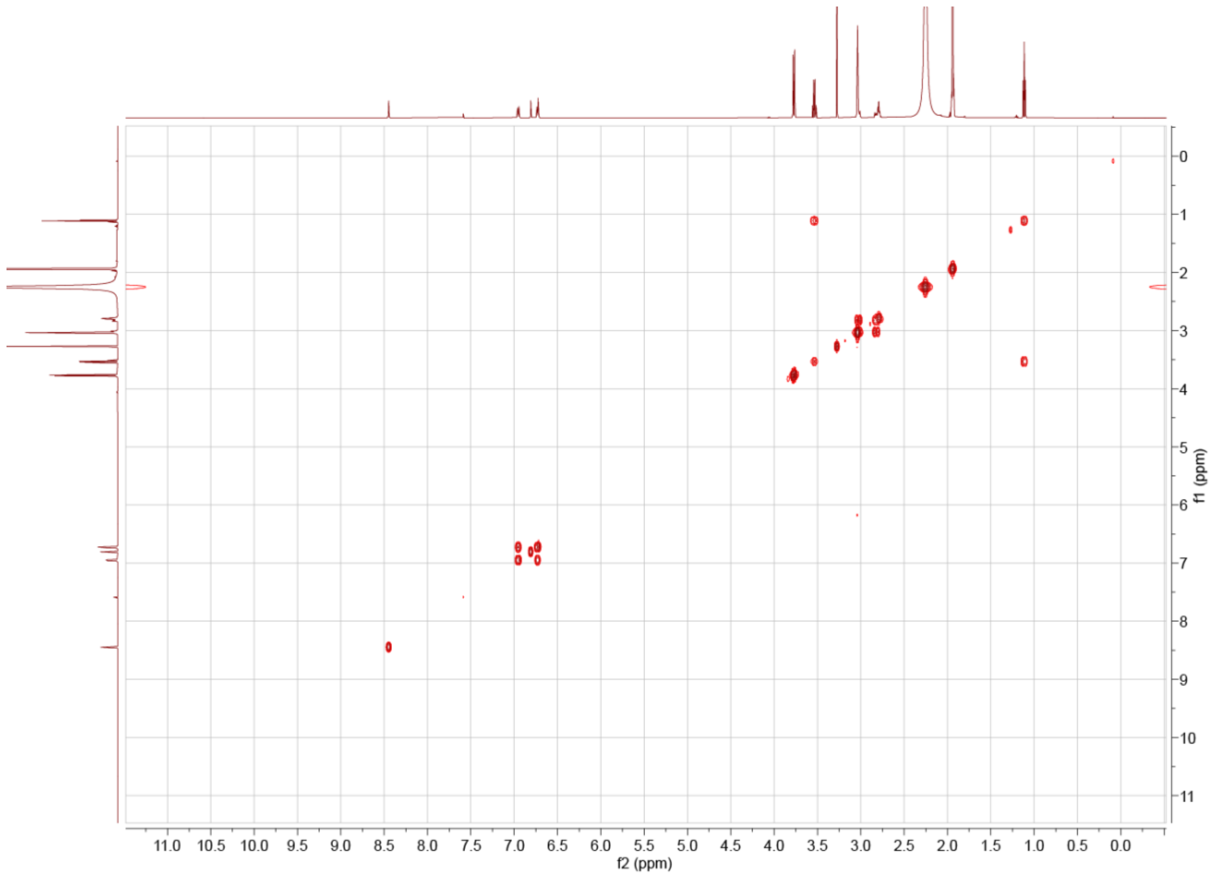


**S. Figure 28**. COSY spectrum of fimbriulatumine (**23**).

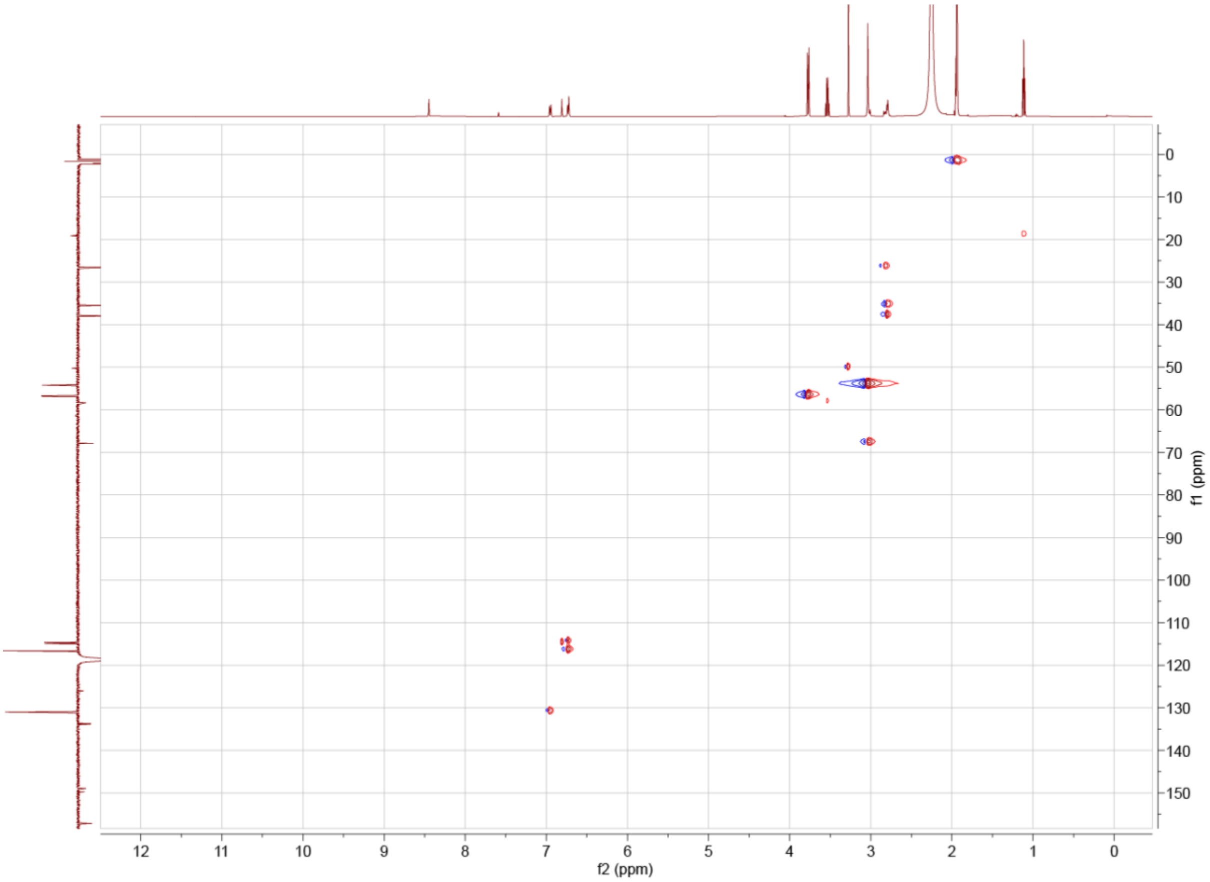


**S. Figure 29**. HSQC spectrum of fimbriulatumine (**23**)

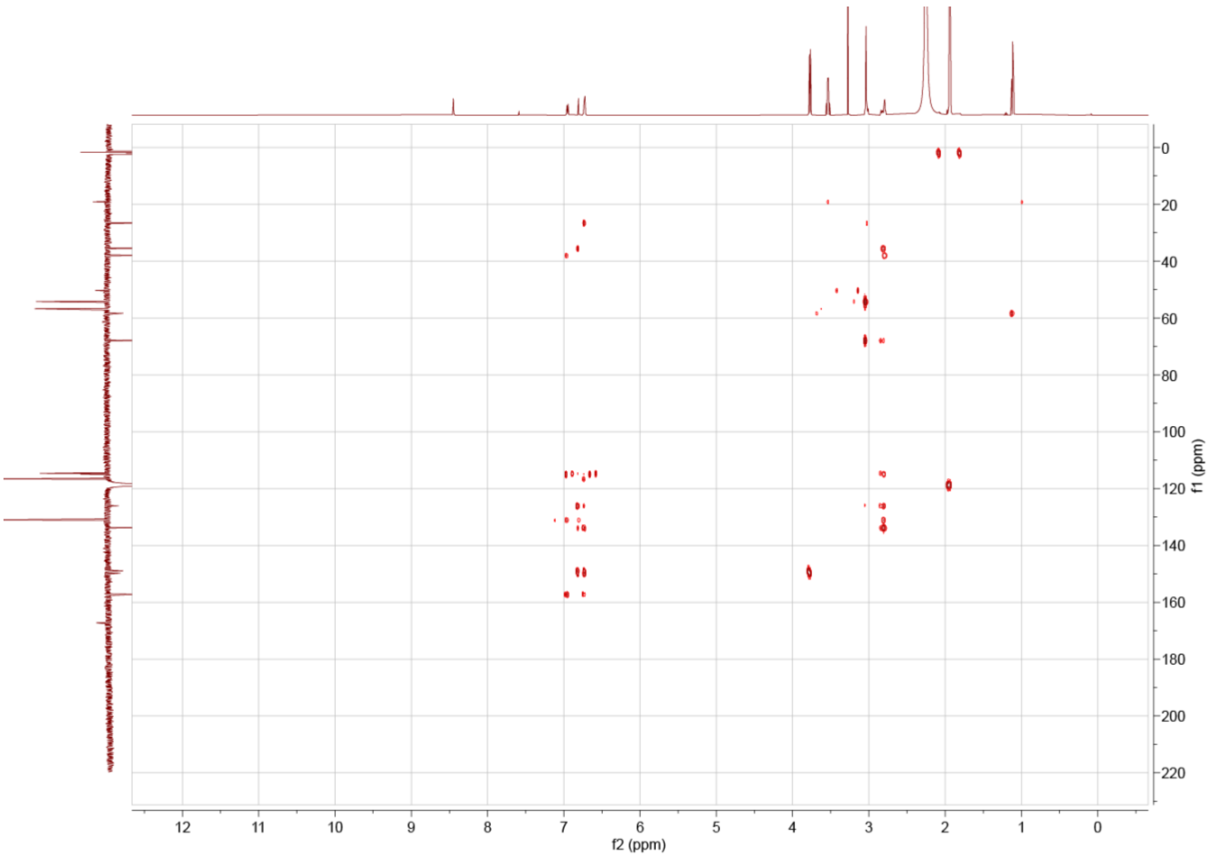


**S. Figure 30**. HMBC spectrum of fimbriulatumine (**23**)

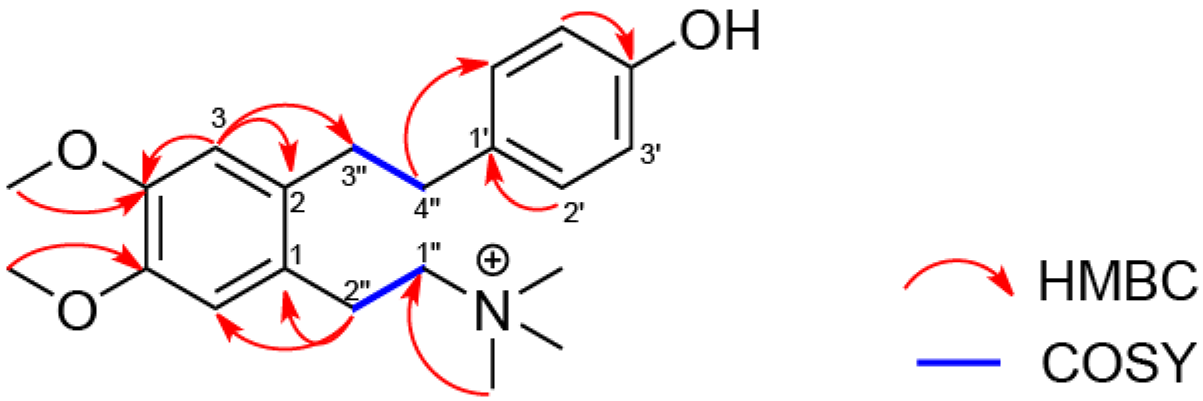


**S. Figure 31**. Structural elucidation of fimbriulatumine (**23**). The $^1$H NMR spectrum showed a sharp singlet ($\delta_H$ = 3.04) integrating to 9 hydrogens indicating a tri-methyl group attached to a nitrogen. This signal ($\delta_H$ = 3.04) couples with C-1'' ($\delta_C$ = 67.5, N–CH2) in HMBC, which is followed by the H-1'' ($\delta_H$ = 3.02) interaction with H-2'' ($\delta_H$ = 2.76–2.84) in COSY and a HMBC C-2'' ($\delta_C$ = 26.2) interaction with H-6 ($\delta_H$ = 6.72) and a H-2'' ($\delta_H$ = 2.76–2.84) interaction with C-1 ($\delta_C$ = 125.7). This connection led us to the linear quaternary ammonium structure connected to an aromatic ring. The doublet of H-3 ($\delta_H$ = 6.81) and H-6 ($\delta_H$ = 6.72) in the $^1$H NMR spectra indicated a benzene ring with 4 connections. Furthermore, the doublet of H-3 ($\delta_H$ = 6.81) and H-6 ($\delta_H$ = 6.72) did not interact with each other in the COSY, thus suggesting they were para. We identified two methoxy groups based on two close singlets ($\delta_H$ = 3.76 and 3.78) with each of them integrating to 3 hydrogens in the $^1$H NMR. HMBC of the methoxy hydrogens ($\delta_H$ = 3.76 and 3.78) are connected to quaternary carbons C-3 and C-4 ($\delta_C$ = 148.6 and 149.4) indicating their connection to a benzene ring. Interactions of the H-3 ($\delta_H$ = 6.81) signal in HMBC confirmed that the hydrogen is sandwiched between C-4 ($\delta_C$ = 149.4) and C-2 ($\delta_C$ = 133.5). The interaction between H-3 ($\delta_H$ = 6.81) and C-2 ($\delta_C$ = 133.5) indicates that the methoxy groups must be next to each other on the same ring and that the molecule continues to another aliphatic group. The last connection to mention is H-3 ($\delta_H$ = 6.81) to C-3'' ($\delta_C$ = 35.1), a different CH2 group than the first. Lastly, the doublet-of-doublet-like signals in the $^1$H NMR spectrum H-2'/6' ($\delta_H$ = 6.95) and H-3'/5' ($\delta_H$ = 6.73) suggested a para substituted ring. HMBC indicates that the H-2'/6' ($\delta_H$ = 6.95) interacts with a CH2, C-4'' ($\delta_C$ = 35.1). Furthermore, HMBC shows that H-2'/6' ($\delta_H$ = 6.95) connects with the quaternary carbon, C-1' ($\delta_C$ = 133.3). This led us to the para substituted ring's connection to the rest of the molecule. Lastly, we see H-3'/5' ($\delta_H$ = 6.73) interact with the quaternary carbon C-4' ($\delta_C$ = 154.8) in HMBC, this carbon shift is indicative of a phenol connected carbon, which explains the doublet-of-doublet-like signals and ultimately gave us fimbriulatumine.

### S. Note 1 - Piperamides annotation

Piperlongumine (**1**) was first putatively annotated via spectral matching against the GNPS MS/MS library (**S. Figure 2A**) and then confirmed using a commercially-available standard (**S. Figure 3A**). Using the MS/MS spectrum obtained from the analytical standard, we identified several diagnostic fragments (i.e., *m/z* 221.081, 206.058, 193.086, 190.063, 178.063, 162.068) generated by the trimethoxycinnamic acid moiety of the molecule, in accordance with previous reports.(da Silva-Junior et al. 2017) Inspection of MN1 revealed three nodes with different precursor mass (i.e., *m/z* 320.150, 306.170, 304.154), but MS/MS spectra nearly identical to piperlongumine (cosine similarity >0.97). The presence of all the trimethoxycinnamic acid diagnostic fragments (**S. Figure 4**) suggests that the modification site is on the piperidine ring. Therefore, we putatively annotated these nodes as dihydropiperlongumine (**2**), 1-(3,4,5-Trimethoxycinnamoyl)piperidine (**3**), 1-(3,4,5-Trimethoxycinnamoyl)-3-piperideine (**4**), respectively. Further inspection of the global molecular network revealed two more potential piperlongumine analogs with precursor mass *m/z* 288.123 and 276.159. Their MS/MS spectra contain several of the diagnostic fragments mentioned above shifted by 30.011 Da ($CH_2O$), thus suggesting a dimethoxycinnamic acid moiety in the molecule (**S. Figure 5**). Therefore, we putatively annotated these compounds as 3'-Demethoxypiperlongumine (**5**) and 1-(3,4-Dimethoxycinnamoyl)piperidine (**6**), respectively. We confirmed our annotation via retention time and MS/MS matching (**S. Figure 3A-E**) using a commercially-available standard for piperlongumine (**1**) and synthetic standards (see **Experimental section**) for compounds (**2**), (**3**), (**5**), and (**6**). In addition, we confirmed the presence of piperine (**8**) in *P. fimbriulatum* roots using a commercially-available standard (**S. Figure 2F**).

Further inspection of MN1 revealed one node with *m/z* 635.260 and retention time 10.41 minutes, which was predicted as piperlongumine dimer (**7**) by CSI:FingerID. Manual inspection of the MS/MS spectrum revealed a high degree of similarity between the spectra (cosine similarity >0.87) and the presence of a fragment peak *m/z* 318.134, most likely corresponding to a loss of piperlongumine monomer (**S. Figure 6**). In order to confirm this annotation, we generated a mixture of piperlongumine dimers by irradiation of the analytical standard with 365 nm UV light (see **Experimental section**) and directly analyzed the reaction mixture using LC-MS. Over the reaction time (from 0 to 30 minutes), a decrease in the amount of the piperlongumine monomer (*m/z* 318.134) was observed concomitantly to an increase in the intensity of all peaks with *m/z* 635.260 (see **S. Figure 7A-B**). The presence of multiple peaks in the extraction ion chromatogram (EIC) of *m/z* 635.260 is likely due to the different regioisomer that can arise from the cycloaddition reaction (i.e., different arrangement of the substituent groups of the cyclobutane ring). Interestingly, the chromatographic profile of the *m/z* 635.260 ion in the reaction mixture overlaps very well with the leaf extract (**S. Figure 7C**). We believe this strongly suggests that the formation of these dimers in the native plant occurs under conditions that are not regio- or stereo-selective, typical of a UV-induced reaction, rather than an enzyme-catalyzed reaction.

#### S. Note 2 - Benzylisoquinoline alkaloids annotation

Higenamine (**9**) was first putatively annotated via spectral matching against the GNPS MS/MS library (**S. Figure 2C**) and then confirmed using a commercially-available standard (**S. Figure 8A**). Similar to what was done for MN1, we used this confirmation to “propagate” the annotation throughout MN2. Using the reference MS/MS spectrum obtained from the analytical standard, we identified several diagnostic fragments (i.e., *m/z* 255.102, 237.091, 219.080, 209.096, 161.060, 149.060, 143.049, 123.044, and 107.049, see **S. Figure 9**), which we used to locate the modification site of potential higenamine analogs within MN2. In particular, MN2 contained three nodes (feature ID 185, 202, 270) with different retention times and same precursor *m/z* 286.144, which likely corresponds to a methylation (+14.015 Da) of higenamine. Manual inspection of the MS/MS spectra of these features strongly supported this hypothesis (**S. Figure 10**). Specifically, we annotated feature 185 (*m/z* 286.144, 3.88 min) as N-methylhigenamine (**12**) due to the presence of the fragment peaks *m/z* 255.102, 178.086, and 161.059 (**$F_1$**, **$F_5$**, **$F_2$**, respectively, in **S. Figure 10**), which suggests the methylation to be on the nitrogen atom. Similarly, we annotated features 202 (*m/z* 286.144, 4.11 min) and 270 (*m/z* 286.144, 5.11 min) as coclaurine (**10**) and its structural isomer isococlaurine (**11**) as they exhibited identical MS/MS spectra (cosine similarity 0.99, mirror MS/MS plot link) and both contained the fragment peaks *m/z* 269.117, 175.075 and 137.059 (**$F_6$**, **$F_7$**, **$F_8$** in **S. Figure 10**), which suggests the methylation to be on either of the two hydroxyl groups in position 4 and 6 of the dopamine moiety. The annotation of coclaurine (**10**) was later confirmed using a commercial standard (**S. Figure 8B**). MN2 also contained six nodes (feature ID 123, 153, 196, 204, 254, 266) with different retention times and same precursor *m/z* 300.160, which likely corresponds to a dimethylation (+28.031 Da) of higenamine. We tentatively assigned the methylation sites based on the manual interpretation of the corresponding MS/MS spectra (**S. Figure 11**). Specifically, we annotated feature 123 (*m/z* 300.160, 3.11 min) as N-dimethylhigenamine (**13**) due to the presence of a dominant peak *m/z* 58.065 (**$F_2$** in **S. Figure 11**), which likely corresponds to the dimethyl(methylene)ammonium fragment (PubChem ID 2724134), and the fragments *m/z* 255.102, 161.060 and 107.049 in the spectrum (**$F_1$**, **$F_3$**, **$F_4$** in **S. Figure 11**). The presence of the *m/z* 58.065 fragment as base peak in MS/MS spectra of BIAs carrying a quaternary nitrogen was later confirmed with commercial standards of compounds (**15**), (**16**) and (**20**) (see below). In contrast, feature 254 (4.91 min) did not contain the peak *m/z* 58.065, but exhibits the fragment peaks *m/z* 283.133, 189.091 and 151.075 (**$F_6$**, **$F_7$**, **$F_8$** in **S. Figure 11**), which indicate two methylations on the two hydroxyl groups in position 4 and 6 of the dopamine moiety. Therefore, we annotated feature 254 as norarmepavine (**14**). Notably, the same *m/z* 58.065 fragments can be also observed in the MS/MS spectrum of armepavine (PubChem ID 442169) which further supports our annotation. A mirror MS/MS plot of feature 254 and the GNPS MS/MS spectrum of armepavine can be generated using the Metabolomics Spectrum Resolver at this link. Concerning features 196, 266, and 204, they all exhibit nearly-identical MS/MS spectra (cosine similarity >0.97) and contain the fragment peak *m/z* 269.117, which suggests one methylation on the nitrogen and a second methylation on one of the three OH groups. This is also supported by the absence of the fragment *m/z* 58.065 as base peak in the MS/MS spectrum, which would originate from a dimethylation on the nitrogen atom. MN2 also contained two features (feature ID 161, 186) with different retention times and same precursor *m/z* 314.176, which likely corresponds to a trimethylation

(+42.046 Da) of higenamine. Both these features exhibit the dominant fragment *m/z* 58.065 ($F_2$ in **S. Figure 11**) and the fragments *m/z* 269.117 and 175.075 ($F_9$ and $F_{10}$ in **S. Figure 11**) which, altogether, suggests a dimethylation on the nitrogen and a third methylation on one of the three OH groups. We confirmed these compounds to be lotusine (**15**) and magnocuranine (**16**), respectively, using commercial standards (**S. Figure 8C-D**). Notably, we observed the base peak *m/z* 58.065 in the MS/MS of both these compounds, which confirms that this fragment arises from the dimethyl(methylene)ammonium fragment upon dimethylation of the nitrogen atom in the BIAs scaffold. Finally, MN2 contained one node (feature ID 227) with *m/z* 328.191, which likely corresponds to a tetramethylation (+56.062 Da) of higenamine. We annotated this feature as N-methylarmepavine (**17**), due to the presence of the base peak *m/z* 58.065, which indicates a dimethylation on the nitrogen atom, and the fragments *m/z* 283.133, 189.091 and 151.075, which indicate two more methylations on the hydroxyl groups of the dopamine moiety.

#### S. Note 3 - Aporphine alkaloids and piperolactams annotation

Asimilobine (**18**), magnoflorine (**20**), and piperolactam A (**21**) were first putatively annotated via spectral matching against the GNPS MS/MS library (**S. Figure 2E-G**). We were particularly confident in the annotation of (**20**) given the presence of the fragment *m/z* 58.065 as base peak in the experimental MS/MS spectrum (red arrow in **S. Figure 2F**), which was an indicator for a dimethylation on the nitrogen in both (**15**) and (**16**). Therefore, we purchased a commercial standard to confirm our annotation (**S. Figure 8F**) and used it as a starting point to "propagate" the annotation throughout the molecular network. Manual inspection of MN3 revealed a node (feature ID 298) with *m/z* 282.149, retention time 5.84 min and a fragmentation pattern in the MS/MS spectra similar to the one observed in (**20**) (see MS/MS mirror plot). We later confirmed this compound to be lirinidine (**19**) via retention time and MS/MS spectral matching (**S. Figure 8E**) using a commercial standard. Concerning (**18**) and (**21**), unfortunately we were not able to purchase the corresponding commercial standards. Therefore, we assigned the annotation based on manual inspection of the experimental and reference MS/MS spectra. In both cases, we assigned the annotation due to the presence of all diagnostic fragments, and similar fragmentation patterns, in the experimental spectra. Differences in the relative intensities ratios between fragments (therefore the decreased cosine similarity) can be explained by the fact that experimental and library MS/MS spectra were acquired on different MS platforms (Orbitrap and time-of-flight, respectively) which are known to produce different spectra.(Heuckeroth et al. 2024)
